# Positive selection of somatically mutated clones identifies adaptive pathways in metabolic liver disease

**DOI:** 10.1101/2023.03.20.533505

**Authors:** Zixi Wang, Shijia Zhu, Yuemeng Jia, Yunguan Wang, Naoto Kubota, Naoto Fujiwara, Ruth Gordillo, Cheryl Lewis, Min Zhu, Tripti Sharma, Lin Li, Qiyu Zeng, Yu-Hsuan Lin, Meng-Hsiung Hsieh, Purva Gopal, Tao Wang, Matt Hoare, Peter Campbell, Yujin Hoshida, Hao Zhu

## Abstract

Somatic mutations in non-malignant tissues accumulate with age and insult, but whether these mutations are adaptive on the cellular or organismal levels is unclear. To interrogate mutations found in human metabolic disease, we performed lineage tracing in mice harboring somatic mosaicism subjected to non-alcoholic steatohepatitis (NASH). Proof-of-concept studies with mosaic loss of *Mboat7*, a membrane lipid acyltransferase, showed that increased steatosis accelerated clonal disappearance. Next, we induced pooled mosaicism in 63 known NASH genes, allowing us to trace mutant clones side-by-side. This *in vivo* tracing platform, which we coined MOSAICS, selected for mutations that ameliorate lipotoxicity, including mutant genes identified in human NASH. To prioritize new genes, additional screening of 472 candidates identified 23 somatic perturbations that promoted clonal expansion. In validation studies, liver-wide deletion of *Bcl6, Tbx3,* or *Smyd2* resulted in protection against NASH. Selection for clonal fitness in mouse and human livers identifies pathways that regulate metabolic disease.

**Highlights:**

1. Mosaic *Mboat7* mutations that increase lipotoxicity lead to clonal disappearance in NASH.
2. In vivo screening can identify genes that alter hepatocyte fitness in NASH.
3. Mosaic *Gpam* mutations are positively selected due to reduced lipogenesis.
4. In vivo screening of transcription factors and epifactors identified new therapeutic targets in NASH.

## INTRODUCTION

Somatic mutations are common in most individuals, and there is accumulating evidence that mutation burden increases with age and chronic tissue damage (Brunner et al., 2019; Lawson et al., 2020; Martincorena and Campbell, 2015; Zhu et al., 2019). While the identity and abundance of these mutations are becoming increasingly understood through deep sequencing studies, many fundamental questions about the relevance of these mutations remain unanswered. The detection of mutant clone expansion, recurrent mutations, or convergent mutations using sequencing provides correlative evidence for increased clonal fitness. However, such fitness increases can be caused by adaptive or pathogenic mechanisms, and it is uncertain if these ever contribute to organ health or function. Even though most somatically mutated clones are not fated to become cancerous, it is still possible that increased proliferation/survival could be entirely selfish and have no beneficial effects on normal tissue function. Therefore, it is unclear how somatic mutations contribute to organismal aging or disease pathogenesis, and more specifically, whether or not somatic mutations can cause a reversal or adaptation to disease.

Recent evidence from the sequencing of human liver disease samples suggests that mutations could be adaptive. Our previous work indicates that some mutations in cirrhotic livers can result in the proliferative expansion of regenerative clones in the context of chemically induced liver injuries (Zhu et al., 2019), however, it is unclear if these expansion events protect against clinically relevant causes of liver disease. With increasing over-nutrition and obesity, non-alcoholic steatohepatitis (NASH) is rapidly becoming the leading cause of liver disease in the world (Diehl and Day, 2017). The excessive accumulation of lipid droplets in hepatocytes eventually leads to lipotoxicity, cell death, and cirrhosis. NASH is usually conceptualized at the organismal and tissue levels, and little thought has been given to genetic heterogeneity between clones in the liver. In liver tissues from NASH patients, Ng et. al. recently identified recurrent and convergent mutations in genes central to insulin signaling and lipogenesis (Ng et al., 2021). The detection of loss of function mutations in metabolic enzymes that generate hepatic lipids suggest that some somatic mutations can confer increased fitness through a reversal of the driving etiology of disease. While these genetic alterations are presumed to be positively selected by virtue of their ability to counteract disease pathogenesis, functional evidence is still required.

To understand the biological importance of somatic mutations at the cellular, tissue, and organismal levels, we developed mouse models that are capable of replicating a high density of mutations in the context of common liver diseases. In this study, the introduction of perturbations in individual genes essential for driving NASH revealed that decreased lipid accumulation can promote clonal fitness and expansion in NASH. Furthermore, we evaluated the fate of mutant clones in a massively parallel, pooled fashion within normal and NASH livers. Remarkably, *in vivo* screening of NASH candidate genes revealed that the somatic mutations detected in human liver tissues are also the most positively selected in mouse models of fatty liver, but are not selected for in the absence of disease. Mechanistically, these mutations reverse lipotoxic phenotypes to increase the survival of hepatocyte clones.

These findings uncover the biological basis for positive selection of somatic mutations in NASH patient livers. Based on these observations, we reasoned that identifying mutant cells with greater fitness than wild-type (WT) cells within diseased environments might nominate new therapeutic targets. This encouraged us to explore genes beyond those that are known to be somatically mutated by performing additional *in vivo* CRISPR screens for genes that are dysregulated in chronic liver disease. These screens identified genes that when inhibited, promote liver fitness through the mitigation of lipotoxicity. We propose that evolutionary selection in somatically mosaic tissues is a new approach for the identification of adaptive metabolic disease pathways and therapeutic targets.

## RESULTS

### The fate of mosaic mutant *Mboat7* clones in the liver is dependent on diet

Fatty liver disease is usually conceptualized at the organismal level and major genes impacting this disease have been identified through the study of germline variants, but the existence of genetically mutant clones within NASH livers is beginning to be recognized. We generated models of fatty liver disease using a Western Diet (21.1% fat, 41% sucrose, and 1.25% cholesterol by weight) supplemented with a high sugar solution (23.1g/L d-fructose and 18.9 g/L d-glucose), a combination hereafter designated as WD. We asked if mosaic mutations in a gene well known to confer an increased predisposition to NASH could also drive fitness differences and competition between liver cells. We focused on *Membrane-bound O-acyltransferase 7* (*Mboat7*), a gene encoding a phospholipid synthetic enzyme identified through GWAS studies (Buch et al., 2015; Xia et al., 2021). To measure the degree to which liver-wide *Mboat7* deletion promotes fatty liver disease in mouse models, we used a high dose of AAV8-TBG-Cre to generate liver-wide knockout (KO) mice. We observed efficient genomic deletion of the floxed exon (**Figure S1A**) and depletion of the intact *Mboat7* mRNA **(Figure S1B)** one week after AAV8-TBG-Cre. After 1.5 months of WD feeding, whole liver *Mboat7* KO vs. WT mice had a dramatic increase in liver/body weight ratios (Figure 1A, **Figure S1C,D**, and **Table S1**). H&E staining showed prominent macroscopic lipid droplets in pericentral hepatocytes of WD fed *Mboat7* KO livers, while only microscopic lipid droplets were observed in WD fed WT livers (Figure 1B). In the context of WD, *Mboat7* deficiency also led to significantly increased transaminitis (**Figure 1C,D**), liver lipid content (**Figure 1E,F**), and lipidemia (**Figure S1E,F**). This data confirms that the presence of *Mboat7* protects against steatosis at the organ level.

**Figure 1.**
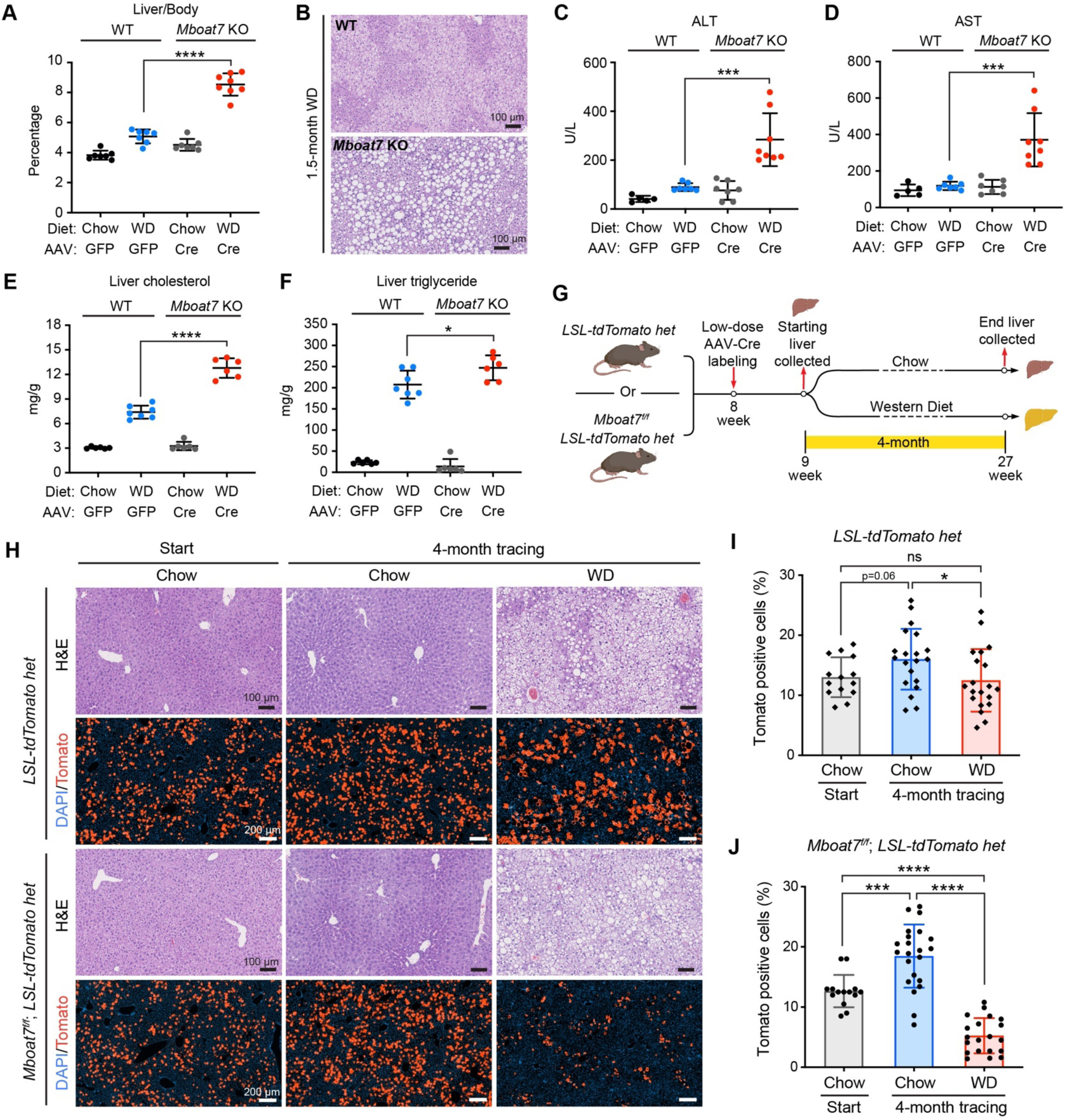
*Mboat7* mutations, which cause excessive lipid accumulation, lead to decreased clonal fitness. A. Liver/body weight ratios of liver-specific *Mboat7* WT and KO mice fed with 1.5 months of chow or WD (n = 7, 7, 7, 8 mice for each group). These mice were given high doses of AAV8-TBG-Cre in order to generate liver-wide *Mboat7* deletion in almost all hepatocytes. B. Representative H&E staining of *Mboat7* WT and KO liver sections after 1.5 months of WD. C-D. Liver function analysis with plasma AST and ALT (n = 5, 7, 7, 8 mice for each group). E-F. Cholesterol and triglyceride measurements from liver tissues (n = 6, 7, 6, 6 mice for each group). G. Schema for the mosaic *Mboat7* lineage tracing experiment. *LSL-tdTomato* het mice or *Mboat7^f/f^; LSL-tdTomato* het mice were injected with a low dose of AAV8-TBG-Cre to generate mosaic Tomato-positive WT hepatocytes in control mice, or Tomato-positive and *Mboat7* mutant hepatocytes in experimental mice. These mice were then fed with either chow or WD for 4 months. Livers were collected one week after AAV8-TBG-Cre, and 4 months after chow or WD was initiated. H. Representative H&E and fluorescent images of liver sections at the beginning and end of lineage tracing. I. Quantification of Tomato-positive cells from *LSL-tdTomato* het liver sections in **H** (n = 7, 10, 10 mice for each group). Each dot represents one image field; two fields from each mouse liver were analyzed. J. Quantification of Tomato-positive cells from *Mboat7^f/f^; LSL-tdTomato* het liver sections in **H** (n = 7, 11, 10 mice for each group). Each dot represents one image field; two fields from each mouse liver were analyzed.

We then wanted to determine if *Mboat7* KO and WT clones have differences in fitness in the context of normal and fatty liver environments. To achieve this, we performed lineage tracing of *Mboat7* WT and KO hepatocyte in mosaic livers. To generate mosaic WT hepatocytes with Tomato reporter activation (control livers) and mosaic *Mboat7* deleted hepatocytes with Tomato reporter activation, we used low dose AAV8-TBG-Cre to achieve random recombination in about 10-15% of hepatocytes in *Mboat7^+/+^*; *Rosa-Lox-stop-Lox-tdTomato* (hereafter *LSL-tdTomato*) and *Mboat7^f/f^*; *LSL-tdTomato* het mice (Figure 1G). Tomato positive clones were traced and quantified under normal chow or WD diets for 4 months (Figure 1H). Body and liver weights increased with WD as expected (**Figure S1G**). The initial Tomato labeling percentages in both groups were similar (**Figure 1I,J left column)**. After 4 months of tracing with chow diets, Tomato positive clones increased in both *Mboat7^+/+^* and *Mboat7^f/f^* mice similarly, (**Figure 1I,J middle column**), indicating a neutral effect of *Mboat7* loss in the liver under normal chow conditions. After 4 months of tracing with WD diets, we observed a modest decrease in the percentage of Tomato positive cells in *Mboat7^+/+^* mice, but a more substantial decrease in *Mboat7^f/f^*mice (**Figure 1I,J right column**). Comparing WT and *Mboat7* KO clones in the WD setting showed that WT clones survived longer and were more fit (**Figure S1H**). When we performed the same lineage tracing experiment over 6 months, the findings were more pronounced, with almost complete disappearance of *Mboat7* KO clones (**Figure S1I-K**). These results demonstrated that somatic mutations in an important lipid enzyme could have a profound impact on clonal survival in a diet and steatosis dependent fashion.

### Analysis of clonal evolution in somatically mutated livers with and without steatosis

These observations supported the concept that mutations conferring fitness differences are subject to strong selection pressures within the fatty liver environment. This also showed that clone size provides a surrogate measure for a gene’s influence on cellular health, not just proliferation, in metabolic liver disease. To expand our understanding of how mutations in different genes might influence clonal dynamics in an unbiased fashion, we developed a platform called MOSAICS (<u>M</u>ethod <u>O</u>f <u>S</u>omatic <u>A</u>AV-transposon In vivo <u>C</u>lonal <u>S</u>creening), which is a CRISPR based method to generate pools of heterogeneous mutant cells within tissues such as the liver. MOSAICS is completely distinct from previous *Fah* KO based regeneration screening systems (Jia et al., 2022; Wuestefeld et al., 2013) because it is designed to assess a much higher density of mutant clones during homeostasis, and does not require rapid proliferation based selection of FAH expressing clones.

While AAVs are optimal for use in the liver, traditional AAVs cannot be used for screening because they do not genomically integrate, and thus their sgRNAs cannot be later quantified. The MOSAICS AAV vector carries a U6 promoter driven sgRNA element and a CAG promoter driven Sleeping Beauty 100 transposase (SB100)-P2A-Cre fusion protein (Figure 2A). The entire AAV payload is flanked by transposon inverted repeat (IR) sequences that enable genome integration of the payload in the presence of SB100 protein (Figure 2A), and thus enables long-term tracing of integrated sgRNAs. Prior studies have combined AAVs with transposons (Cooney et al., 2015; Ye et al., 2019), but components within the MOSAICS vector and AAV were specifically engineered to meet the needs of generating and tracing mutations in the liver. We assessed the dose dependent effects of a MOSAICS vector containing an sgRNA against *Pten*. We injected the MOSAICS AAV8-*sgPten* into doxycycline (dox)-inducible *TetO-Cas9* mice to generate somatic mutations in the liver, or into *LSL-tdTomato* mice to monitor the expression of SB100-P2A-Cre protein, which activates Tomato (Figure 2B). Increasing amounts of AAV could delete *Pten* or activate the Tomato reporter in an increasing number of hepatocytes (**Figure 2C-E**), indicating proper functioning of the vector elements and that this system could be titrated to generate different levels of somatic mosaicism, or alternatively, liver-wide gene deletion.

**Figure 2.**
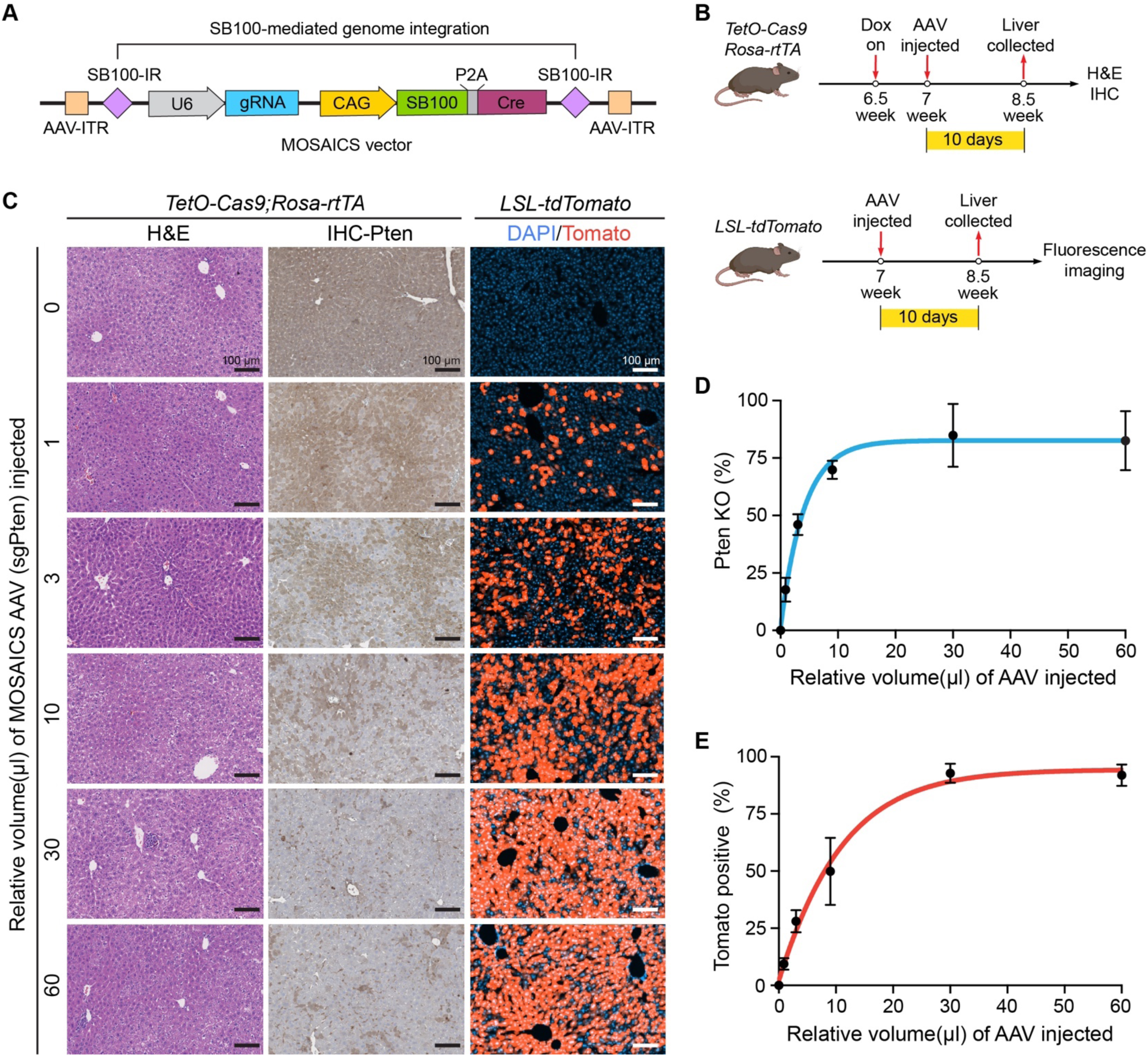
Design of MOSAICS platform for generating and tracing somatic mutations *in vivo*. A. Design of the MOSAICS vector, which integrates AAV delivery, gene perturbation, and sgRNA tracing together. The U6-sgRNA element and CAG promoter-driven SB100-P2A-Cre fusion protein are flanked by inverted repeat (IR) sequences, which enable SB100 transposase mediated genomic integration. B. Schematic for functional validation of the MOSAICS platform. The MOSAICS AAV carrying a *Pten* sgRNA was IV injected into dox-inducible Cas9 expressing mice (*TetO-Cas9* homo; *Rosa-rtTA* homo) to examine the generation of *Pten* mutant hepatocytes in the liver. The same AAV was also injected into *LSL-tdTomato* homo mice to test the expression of the SB100-P2A-Cre fusion protein, which turns on Tomato expression in hepatocytes. C. Liver sections from the validation mice described in B. H&E staining and PTEN IHC staining showed that the frequency of PTEN deficient hepatocytes correlated with the amount of AAV injected. Fluorescent images from liver sections of *LSL-tdTomato* mice showed the frequency of hepatocytes expressing Tomato correlated with the amount of AAV injected. D. Quantification of *Pten* KO cells shown in **C** (n = 3 mice for each AAV concentration). E. Quantification of Tomato positive cells shown in **C** (n = 3 mice for each AAV concentration).

After validation of the MOSAICS system, we next aimed to determine if genes known to be important in NASH would have substantial effects on clonal fitness. We generated an sgRNA library against 63 NASH genes including those identified through somatic mutation sequencing (*GPAM, ACVR2A, FOXO1*), GWAS (*MBOAT7, TM6SF2, GCKR*), exome-seq (*PNPLA3, HSD17B13*), biochemical studies (Samuel and Shulman, 2018), or drug targets (ACC1/2, FXR, ASK1) (**Figure S2A**). We generated mosaically mutated livers by injecting the AAV library into Cas9 expressing mice, and then exposed the mice to either normal chow or WD. Deep sequencing and analysis of enriched or depleted guides after a fixed time period enabled us to monitor clonal dynamics in both chow and WD conditions (Figure 3A). By including sgRNAs specifically enriched in WD conditions and excluding sgRNAs enriched in both chow and WD conditions, we were able to identify fatty liver specific, fitness promoting mutations and exclude genes whose mutations induce a general, fatty liver independent proliferation. Seven to eight independent mice were used for each group, and the results from individual mice were largely consistent (**Table S2**). While liver weight, body weight, and steatosis increased as expected on WD (**Figure S2B,C**), we did not detect surface liver tumors in mice fed with WD for 6 months, suggesting that the somatic mutations did not cause rapid cancer development.

**Figure 3.**
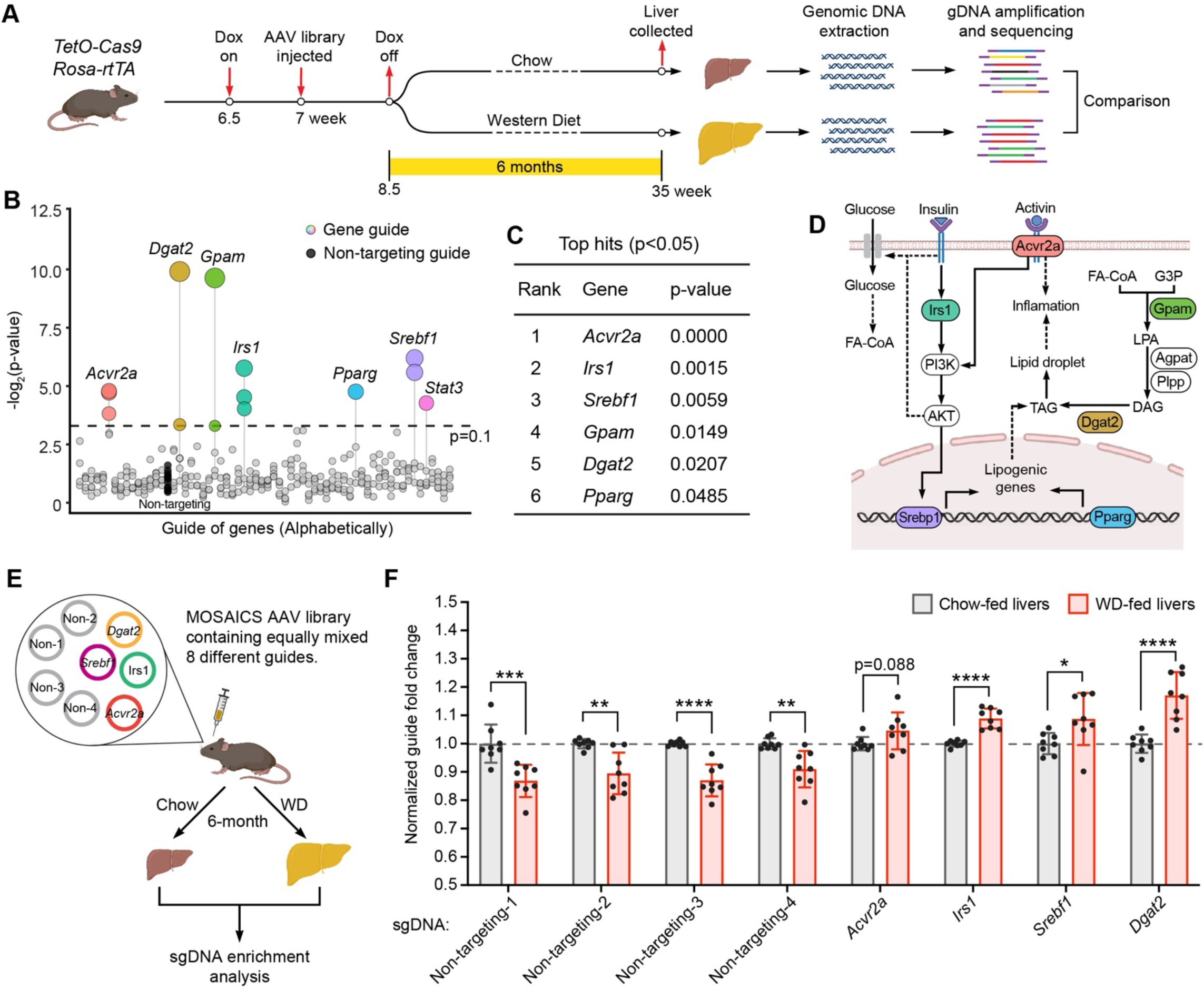
Lineage tracing of mosaic mutant hepatocytes demonstrated that mutations that suppress lipogenesis become enriched in fatty livers. A. Schema for pooled tracing of mosaic mutant hepatocytes under different dietary conditions. MOSAICS AAVs carrying sgRNA libraries were injected into Cas9 expressing mice. Ten days after gene perturbation, Cas9 expression was turned off by dox withdrawal, and chow or WD was given to mice for another 6 months. Genomic DNA was extracted from harvested livers. sgRNA sequences were amplified and quantified. sgRNAs that were specifically enriched in fatty but not normal livers were further investigated. This approach aimed to exclude sgRNAs that target proliferation suppressing genes which would manifest as enrichment in both normal and fatty livers. B. sgRNAs enriched in WD fed but not chow fed livers. Each circle represents one sgRNA. Different sgRNAs targeting the same genes were aligned vertically. Circle sizes correlate to -log2(p). Control sgRNAs were drawn as filled black circles. See **Table S2** for raw data. C. Genes associated with sgRNAs enriched in WD fed mice (p < 0.05). See Table S2 for raw data. D. Pathways in which the enriched genes (listed in **C**) are involved. E. Validation of the MOSAICS platform for the most enriched mutations in fatty livers. Four distinct non-targeting sgRNAs and the sgRNAs targeting *Acvr2a*, *Irs1*, *Srebf1* and *Dgat2* (Table S5) were cloned into the MOSAICS vector. The eight vectors were mixed equally before being used for AAV production. Cas9 mice were injected with the 8-sgRNA AAV library and treated with chow or WD for 6 months. Genomic DNA was extracted from collected livers and sgRNA sequences were amplified by PCR and quantified by high-throughput sequencing. F. Normalized sgRNA reads in chow or WD fed livers (n = 8 and 8 mice for each group). See Table S2 for raw data.

The five genes associated with the highest levels of clonal expansion in the WD but not chow group were *Acvr2a*, *Gpam*, *Dgat2*, *Srebf1*, and *Irs1* (**Figure 3B,C** and **Table S2**). Each is well known to be important in NASH (Figure 3D). *Dgat2* and *Gpam* are critical enzymes in triglyceride synthesis, and for this reason, *Dgat2* is a therapeutic target currently being tested in NASH clinical trials (Calle et al., 2021; Yenilmez et al., 2022). *Srebf1* is a major transcriptional regulator of lipogenesis (Briggs et al., 1993; Brown and Goldstein, 1997; Wang et al., 1993). *Irs1* is an insulin receptor substrate and when lost, leads to suppressed lipogenesis due to reduced insulin signaling (Sun et al., 1991). Interestingly, *Acvr2a* and *Gpam* are two of the most recurrently mutated genes in cirrhotic tissues from alcoholic liver disease and NASH patients (Ng et al., 2021). *Foxo1* gain of function hotspot mutations were also recurrently identified as clonally expanded in human tissues; consistently, *Foxo1* loss of function mutations were found to be strongly associated with clonal disappearance in the MOSAICS screen (**Table S2**). To further assess the screening results with a smaller set of genes that allowed for deeper sgRNA coverage, we fate mapped a mini-pool of sgRNAs against *Acvr2a*, *Irs1*, *Srebf1*, and *Dgat2* as well as 4 non-targeting sgRNAs (Figure 3E). After 6 months, the control sgRNAs in the WD fed livers consistently decreased in abundance compared to those in chow fed livers, whereas each of the guides against *Acvr2a*, *Irs1*, *Srebf1*, and *Dgat2* were positively selected (Figure 3F and **Table S2**). The depletion of control guides was likely due to the increased turnover of hepatocytes in WD fed conditions, as was also seen for Tomato in Figure 1I (right vs. middle column). These data show that multiple mutations that impair lipid accumulation result in a fitness advantage for clones in the NASH environment.

### *Gpam* mutant clones are more fit and exert a beneficial effect on the liver

We aimed to perform a deeper investigation of mutations in *Gpam*, which was a top hit from the functional NASH screen described above and from human somatic mutation sequencing efforts (Ng et al., 2021). GPAM catalyzes the acylation of glycerol-3-phosphate with acyl-coenzyme A (CoA) to generate CoA and lysophosphatidic acid (LPA), which is the rate-limiting step in triacylglycerol synthesis (Figure 3D). However, no one has studied mosaic *Gpam* mutant clones in comparison with WT clones in the fatty liver microenvironment. Because the conditional KO mice required to study liver specific deletion or to visualize clonal dynamics were not available, we used germline CRISPR approaches to generate *Gpam* floxed mice targeting exon 3, whose deletion is predicted to lead to a frameshift and a premature stop codon. We confirmed that saturating doses of AAV8-TBG-Cre in *Gpam* floxed mice led to the deletion of exon 3 and almost complete depletion of *Gpam* mRNA one week after injection (**Figure S3A,B**). Using these floxed mice, we asked how mosaic WT and KO clones in the liver would compare under normal and WD conditions. In similar fashion as the *Mboat7* experiments, we used low dose AAV8-TBG-Cre to induce Tomato in 10-15% of hepatocytes in *LSL-tdTomato* het mice (control group), or to induce Tomato with *Gpam* deletion in 10-15% of hepatocytes in *Gpam^f/f^; LSL-tdTomato* het mice, then gave these mosaic mice either chow or WD (**Figure 4A,B**). After 6 months of WD, the expected increases in body and liver weights were observed (**Figure S3C**). Although there were similar initial Tomato labeling frequencies in both groups (**Figure 4C,D left column**), and a similar level of *Gpam* WT and KO clonal expansion in the setting of chow (**Figure 4C,D middle column**), we observed an increase in *Gpam* KO clone abundance compared to WT in the setting of WD (**Figure 4C,D right column**). These data demonstrate the diet and steatosis context dependent effects of *Gpam* mutations, which is also in accord with the *Mboat7* tracing experiments (**Figure 1H-J**).

**Figure 4.**
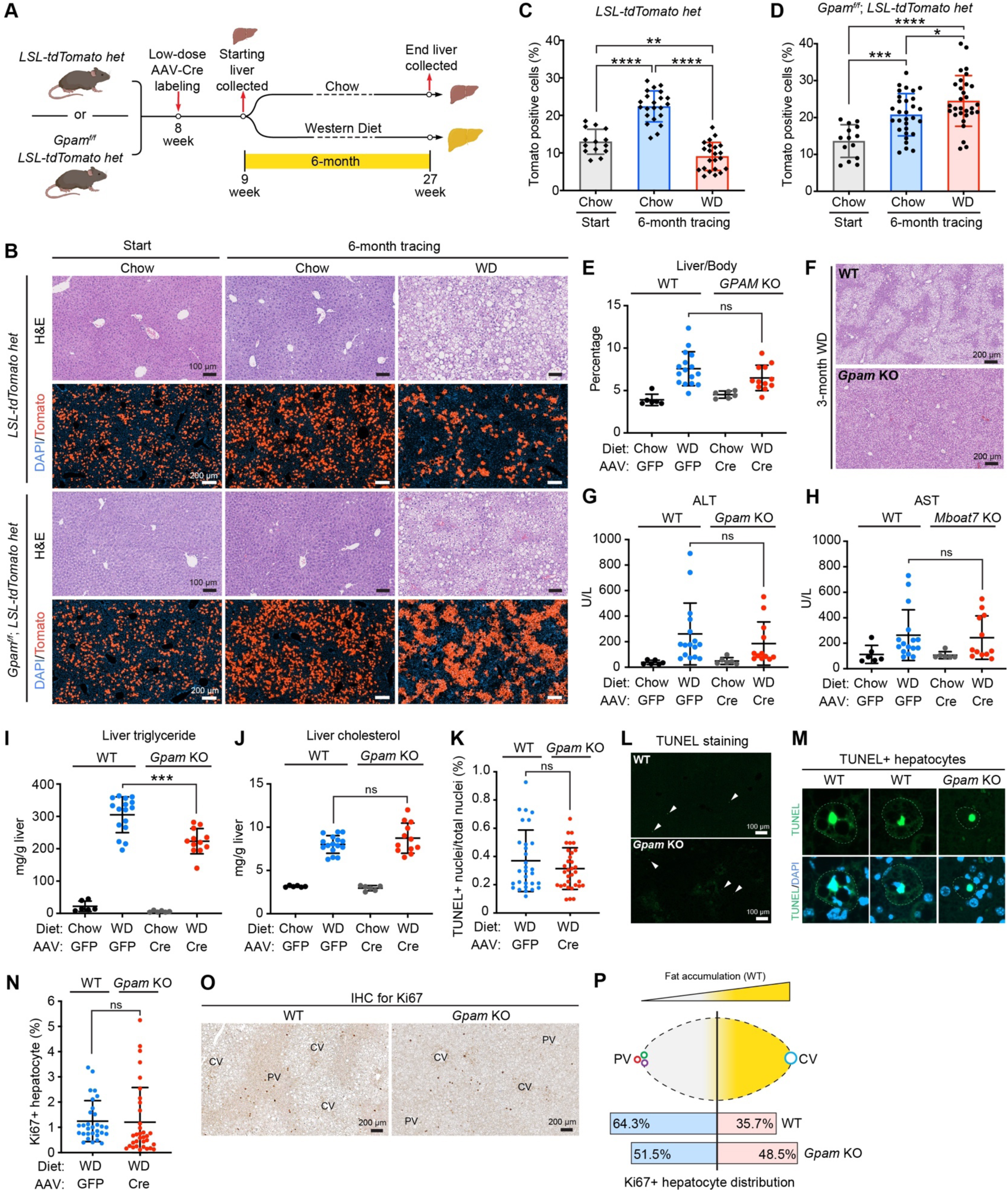
*Gpam* mutations, which suppress lipogenesis, lead to increased clonal fitness. A. Schema for the mosaic *Gpam* lineage tracing experiment. *LSL-tdTomato* het mice or *Gpam^f/f^; LSL-tdTomato* het mice were injected with a low dose of AAV8-TBG-Cre to generate mosaic Tomato-positive hepatocytes in control mice, or Tomato-positive and *Gpam* mutant hepatocytes in experimental mice. These mice were then fed with either chow or WD for 6 months. Livers were collected one week after AAV injection, and 6 months after chow or WD was initiated. B. Representative H&E and fluorescent images of liver sections at the beginning and end of lineage tracing. C. Quantification of Tomato-positive cells from *LSL-tdTomato* het liver sections in **B** (n = 7, 11, 11 mice for each group). Each dot represents one image field; two fields from each mouse liver were analyzed. The same time zero group of mice was used in Figure 1I. D. Quantification of Tomato-positive cells from *Gpam^f/f^; LSL-tdTomato* het liver sections in **B** (n = 7, 15, 15 mice for each group). Each dot represents one image field; two fields from each mouse liver were analyzed. E. Liver/body weight ratios of liver-specific *Gpam* WT and KO mice fed with 3 months of WD (n = 6, 16, 6, 12 mice for each group). These mice were given high doses of AAV8-TBG-Cre in order to generate liver-wide *Gpam* deletion in almost all hepatocytes. F. Representative H&E staining of *Gpam* WT and KO liver sections after 3 months of WD. G-H. Liver function analysis with plasma ALT and AST (n = 6, 16, 6, 12 mice for each group). I-J. Triglyceride and cholesterol measurements from liver tissues (n = 6, 16, 6, 12 for each group) K. Quantification of TUNEL positive cells from liver sections of *Gpam* WT and KO mice fed with 3 months of WD (n=10 and 11). Each dot represents one image field; three fields from each mouse liver were analyzed. L. Representative TUNEL staining of *Gpam* WT and KO liver sections after 3 months of WD. M. Representative TUNEL positive hepatocytes from *Gpam* WT and KO liver sections after 3 months of WD. N. Quantification of Ki67 positive hepatocytes from liver sections of *Gpam* WT and KO mice fed with 3 months of WD (n = 10 and 11). Each dot represents one image field; three fields from each mouse liver were analyzed. O. Representative Ki67 IHC staining of *Gpam* WT and KO liver sections after 3 months of WD. P. Distribution of Ki67 positive hepatocytes in the portal vein (PV) half and central vein (CV) half of the lobule.

To explore the mechanistic basis for the fitness increases of *Gpam* mutant clones, we investigated the whole liver *Gpam* KO mice induced with AAV8-TBG-Cre. Although no statistically significant body and liver weight differences were observed between WT and whole-liver *Gpam* KO mice after 3 months of WD (**Figure S3D,E** and **Table S1**), the KO group did show a trend toward decreased liver/body ratios (Figure 4E), significantly decreased steatosis (Figure 4F), and a trend toward improved transaminitis (**Figure 4G,H**) on WD. Liver triglyceride but not cholesterol levels were significantly decreased in the WD fed KO group (**Figure 4I,J**), consistent with *Gpam*’s role as a rate limiting enzyme in triglyceride synthesis. Interestingly, plasma cholesterol but not triglyceride was decreased in the KO group (**Figure S3F,G**). We attempted to determine if increased survival or proliferation were responsible for the clonal expansion of *Gpam* deficient hepatocytes in the WD setting. The frequency of TUNEL positive nuclei in WT and *Gpam* KO livers were similar and low (**Figure 4K,L**). In WT livers, some apoptotic hepatocytes were filled with macroscopic lipid droplets, whereas in KO livers, the apoptotic hepatocytes did not have these characteristics (Figure 4M). Similarly, WT and KO liver sections showed comparable frequencies of proliferating hepatocytes as measured by Ki67 (**Figure 4N,O**), suggesting that liver-wide *Gpam* KO does not lead to overproliferation. However, proliferating hepatocytes in WT livers were more likely to be found near portal triads, where steatosis is less pronounced, whereas proliferating hepatocytes in KO livers were more evenly distributed between portal and central regions (**Figure 4O,P**). This suggests that steatotic hepatocytes are less likely to divide, and that the small proliferative advantage of *Gpam* KO over WT cells could result in expansion under steatotic conditions over long time periods (Figure 4B). Altogether, our data and previous studies show that *Gpam* loss of function mutations lead to fitness-promoting clonal expansions in NASH, and that *Gpam* is a promising therapeutic target in NASH.

### Somatic mosaic screening of candidate genes identifies putative targets for NASH

Positive selection of specific clones in the NASH environment suggests that some somatic mutations can promote fitness through altered lipid accumulation or resistance to lipotoxicity. Therefore, clone size/sgRNA abundance can be a surrogate measure for a gene’s influence on cellular fitness in NASH livers. Based on fitness competition, MOSAICS represents an entirely new way of identifying NASH genes. This concept was strengthened by the fact that the same mutated genes such as *Gpam* and *Acvr2a* were selected for in both mouse and human tissues (Ng et al., 2021). Equipped with this novel method, we sought to discover new functionally important genes and fitness-promoting mutations in NASH using MOSAICS. We reasoned that transcription or epigenetic factors might have the strongest impact on a broad array of pathways, so we analyzed human gene expression data to nominate candidate factors whose activities are either increased or decreased chronic liver disease (**Figure S4**). We performed modeling of putative transcription factor activities using gene expression data from NASH (72 patients; GSE130970) (Hoang et al., 2019) and HCV cirrhosis cohorts (216 patients; GSE15654) (Hoshida et al., 2013). Since transcription factor gene expression levels do not always reflect their functional importance, we chose those with the highest or lowest activities based on the induction or suppression of their downstream targets (See methods for details) (Liberzon et al., 2011; Subramanian et al., 2005). In contrast, the epigenetic genes were ranked only based on their differential expression in human cirrhosis samples. To further narrow down the most influential factors, candidate genes were also associated with histologic features such as fibrosis, inflammation, ballooning, and steatosis, as well as temporal events such as Child-Pugh score, liver decompensation, cancer development, and death. Using this clinico-genomic pipeline, we identified 217 transcriptional and 255 epigenetic regulators that have the highest likelihood of having an impact in human NASH (**Table S3**).

We next created sgRNA libraries corresponding to these two gene sets (transcription and epigenetic factors), and used them to generate somatically mutated mouse livers using the MOSAICS platform (Figure 3A and **Figure S5**). After 6 months of chow or WD, the sgRNAs associated with the most clonal expansion in WD livers targeted 13 and 10 genes in each screen, respectively (**Figure 5A-C** and **Table S2**). Several genes were previously connected with fatty liver disease such as the glucocorticoid receptor *Nr3c1* (Moreira et al., 2014; Wang et al., 2020), the mineralocorticoid receptor *Nr3c2* (Pizarro et al., 2015; Wada et al., 2013), the transcription factor *Zbtb16* (Liška et al., 2017), the DNA methyltransferase *Dnmt1* (Kutay et al., 2012), and the E3 ubiquitin ligase adaptor *Keap1* (Mohs et al., 2021; Ramadori et al., 2016), but most of the top genes had no known connection with liver disease or NASH.

**Figure 5.**
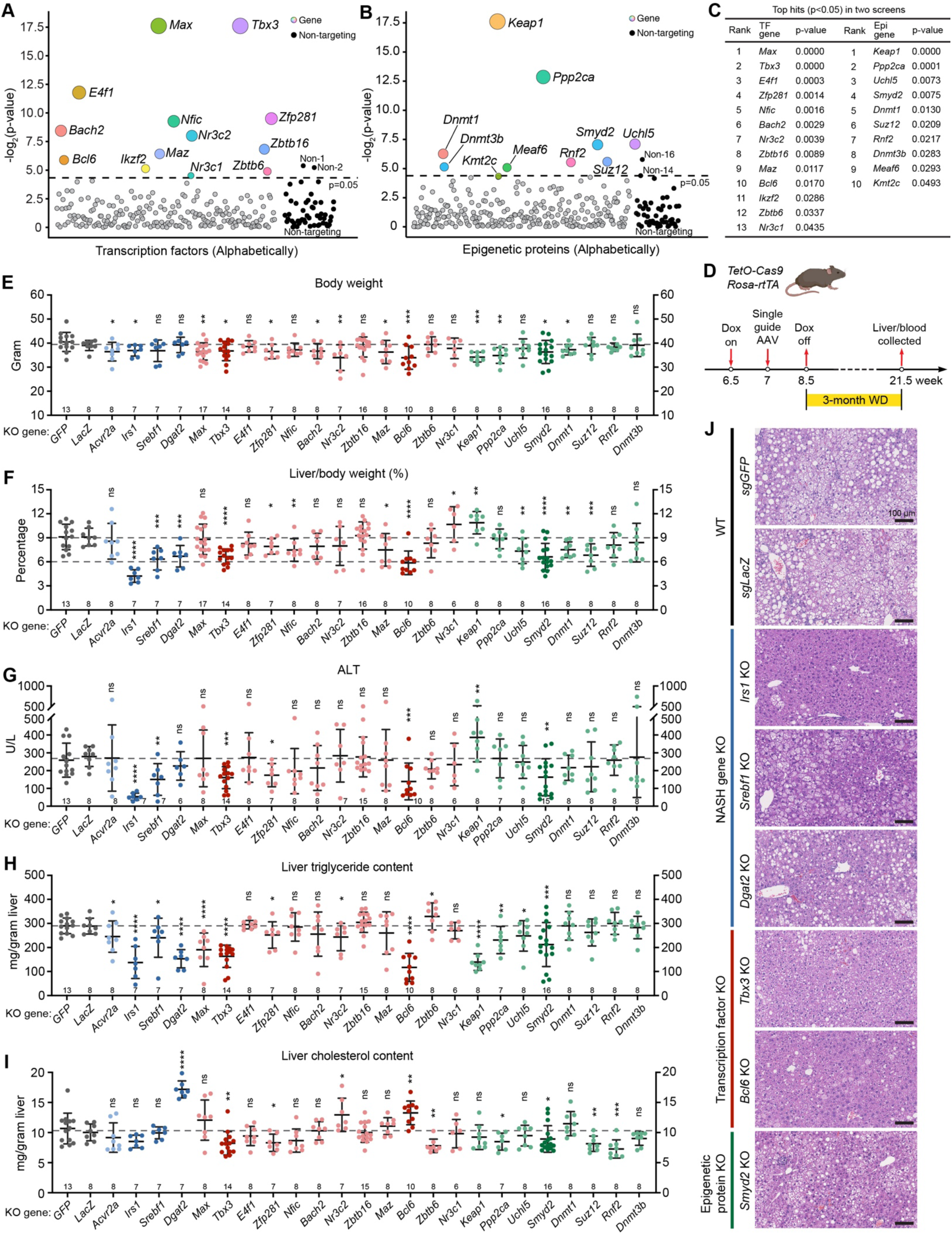
Somatic mosaic screening of transcription and epigenetic factors identifies putative therapeutic targets for NASH. A. Results for somatic mosaic screening of transcription factors. MOSAICS vectors carrying transcription factor targeting sgRNA libraries were injected into Cas9 expressing mice. Chow or WD was fed to mice for 6 months. The genes corresponding to enriched sgRNAs in WD fed but not chow fed livers were drawn as colored circles with sizes correlating to -log2(p). Control sgRNAs were drawn as filled black circles. B. Results for somatic mosaic screening of epigenetic proteins. The screening method and color scheme are the same as in **A**. C. List of the genes corresponding to enriched sgRNAs (p < 0.05) in both MOSAICS screens. D. Schematic for examining the KO phenotypes of the top genes under WD conditions. A MOSAICS AAV carrying an individual sgRNA was injected into Cas9 mice such that each mouse had one gene deleted in the liver. Dox was withdrawn 10 days after AAV injection and WD was given for 3 months before sacrifice. E-F. Body weight and liver/body ratios of control (sgGFP and sgLacZ) and liver-specific KO mice fed with 3 months of WD. Gray dots represent control mice, blue dots represent liver-specific KO mice for known NASH genes, red dots represent liver-specific transcription factor KO mice, and green dots represent liver-specific epigenetic factor KO mice. Darker dots represent mice that have the most significant differences in liver/body weight ratios. Each dot represents one mouse, and the n is denoted at the bottom of each plot. G. Liver function analysis using plasma ALT. The color scheme is the same as in **E**. The n is denoted at the bottom of the plot. H-I. Liver triglyceride and cholesterol analysis. J. Representative H&E images of liver sections are shown for the mice described in **E**.

### Establishment of conditional deletion methods for rapid liver disease phenotyping

To further ascertain which of the top genes, when genetically ablated in all hepatocytes, have the most positive influences on metabolism under WD conditions, we developed rapid conditional KO approaches. This also allowed us to compare the extent of lipid reduction between these top gene mutations and with *bona fide* NASH regulators identified above (*Acvr2a, Irs1, Srebf1, Dgat2*). We cloned the most enriched sgRNA for each of the top genes into the MOSAICS vector, then delivered saturating doses of these AAVs with individual guides into Cas9 expressing mice to achieve single-gene whole-liver deletion (as shown in **Figure 2C,D**). We first tested this system using the four known NASH genes that scored as the most positively selected hits from the 63 NASH gene screen (**Figure 3**). High titers of MOSAICS AAV8-sgRNAs against *GFP* (control 1), *LacZ* (control 2), *Acvr2a, Irs1, Srebf1* or *Dgat2* (**Figure S6A**) were injected into Cas9 expressing mice at 6.5 weeks of age, then were fed WD for 3 months (Figure 5D). Prior to WD feeding, we assessed liver Cas9 cutting efficiency using qPCR based genotyping (**Figure S6B**). The remaining intact mRNAs of the targeted genes in KO livers fell to 25-50% of the levels in WT livers, indicating efficient somatic gene editing (**Figure S6B**). After 3 months of WD, each conditional CRISPR KO mouse was phenotyped for body weight, liver weight, liver lipids (triglyceride, cholesterol), plasma parameters (ALT, AST, triglyceride, cholesterol) and histology (**Table S1**). There were significant reductions in liver to body weight ratios for *Irs1*, *Srebf1*, and *Dgat2* liver KO mice (**Figure 5E,F** and **Figure S6C**). Furthermore, the protection against fatty liver disease was evident from improved plasma parameters, decreased liver lipid content (**Figure 5G-I** and **Figure S6D-F**), and reduced steatosis (Figure 5J and **Figure S7A,B**). These findings were tightly correlated with the degree of liver/body weight ratio reduction, a surrogate marker of fat accumulation and NASH. These data showed that CRISPR mediated liver-wide conditional deletion would be an effective and rapid method of organ level validation.

### Mutations in *Tbx3*, *Bcl6*, and *Smyd2* showed protection against fatty liver disease

An advantage of this rapid liver KO system was that it allowed us to rank the impact of the top genes in the transcription and epigenetic factor screens, and to compare these novel genes with canonical NASH genes. Among all 20 KO mice associated with the transcription and epigenetic factor screens (**Table S1**), more than half of the KO models showed significant reductions in liver mass and none showed significant increases (**Figure S6C**). This suggested that mutations that drive clonal expansions in WD fed livers more often led to reduced lipid accumulation in the liver. Among all KO models, deletion of the transcription factors *Tbx3* and *Bcl6*, as well as the epigenetic factor *Smyd2*, showed significant reductions in body weights (Figure 5E) and the most substantial reductions in liver/body weight ratios compared to controls after 3 months of WD (dark dots in Figure 5F). *Bcl6* is a transcriptional repressor that is enriched in the fed state and cooperates with PPARα to suppress the induction of fasting transcription, and *Bcl6* loss has been shown to increase insulin sensitivity and repress NASH (Chikada et al., 2020; Senagolage et al., 2018; Sommars et al., 2019). Neither *TBX3*, a T-box transcription factor (Willmer et al., 2017), nor *SMYD2*, a protein lysine methyltransferase with histone and non-histone methylation targets, has been studied in fatty liver disease. These three KO models showed the most significant reductions in liver injury as measured by ALT (Figure 5G), a trend toward reduced AST (**Figure S6D**), and the most significant reductions in liver triglyceride levels (Figure 5H). While *Tbx3* and *Smyd2* KO mice had reduced liver cholesterol, *Bcl6* KO mice had increased liver cholesterol levels, which is similar to that of *Dgat2* KO mice (Figure 5I). We also observed reduced plasma cholesterol in *Smyd2* KO mice and increased plasma triglyceride in *Tbx3* KO mice, while the other KO models showed modest or no changes in these parameters (**Figure S6E,F**). Importantly, we observed a clear reduction in hepatic steatosis in *Tbx3, Bcl6*, and *Smyd2* KO livers compared with controls and other KO models (Figure 5J and **Figure S7C,D**). The phenotypes of *Tbx3, Bcl6*, and *Smyd2* KO mice were comparable to *Srebf1* and *Dgat2* KO mice, suggesting potent regulation of lipid metabolism by these three genes.

### Transcriptional analysis revealed diverse mechanisms of lipid regulation

To investigate the gene expression changes in WD treated livers carrying fitness promoting mutations, we performed RNA-seq on control (sgGFP, sgLacZ), *Irs1, Srebf1, Tbx3, Bcl6,* and *Smyd2* KO livers that were generated with saturating doses of MOSAICS AAVs (**Table S4**). We observed massive expression changes in *Irs1* KO livers and fewer changes in *Tbx3* and *Bcl6* KO livers (Figure 6A), which is congruent with *Irs1* KO phenotypes showing the most substantial decreases in body weight, liver weight, inflammation, and steatosis, and supports the known regulation of lipogenesis by insulin signaling (**Figure 5E-J**). *Srebf1* KO livers showed fewer gene expression changes compared with *Irs1*, *Tbx3*, and *Bcl6* KO livers; *Smyd2* KO livers showed the smallest number of genes with expression changes (Figure 6A). Comparing the differentially regulated genes in control vs. KO livers showed a relatively large intersection between *Tbx3* and *Irs1* KO livers (Figure 6B), suggesting shared regulatory circuits. The intersections among *Bcl6*, *Tbx3*, and *Srebf1* KO livers were smaller (Figure 6C), suggesting unique transcriptional roles. Using Gene Set Enrichment Analysis (GSEA), we observed reduced inflammatory pathways in *Irs1, Srebf1*, and *Tbx3* KO livers and reduced proliferation in all KO compared with control livers (Figure 6D), indicating decreased cell turnover in KO livers. For lipid metabolism pathways in GSEA analysis, we observed some shared but mostly unique expression patterns for each KO group (Figure 6D). We further examined the expression of genes involved in lipogenesis and fatty acid oxidation. Deletion of *Irs1*, *Srebf1*, *Tbx3*, *Bcl6*, and *Smyd2* each led to decreased expression of fatty acid and triglyceride synthesis genes, but to different extents (Figure 6E). We also observed decreases in cholesterol synthesis genes in *Irs1* and *Smyd2* KO livers, but a dramatic upregulation of these genes in *Srebf1* KO livers (Figure 6E). Since *Srebp1* is a master transcription factor for cholesterol synthesis, this increase was potentially due to a compensatory effect on cholesterol production. For lipid degradation pathways, strong decreases in β-oxidation genes were observed in *Irs1* and *Bcl6* KO livers (Figure 6F **upper panel**), likely because these livers have the least triglyceride content among all groups (Figure 5H). We observed upregulation of many TCA cycle genes in *Tbx3* and *Smyd2* KO livers (Figure 6F **lower panel**), suggesting that these livers have higher rates of acetyl-CoA consumption. These data indicate shared and unique mechanisms by which mutant hepatocytes converge on decreased steatosis.

**Figure 6.**
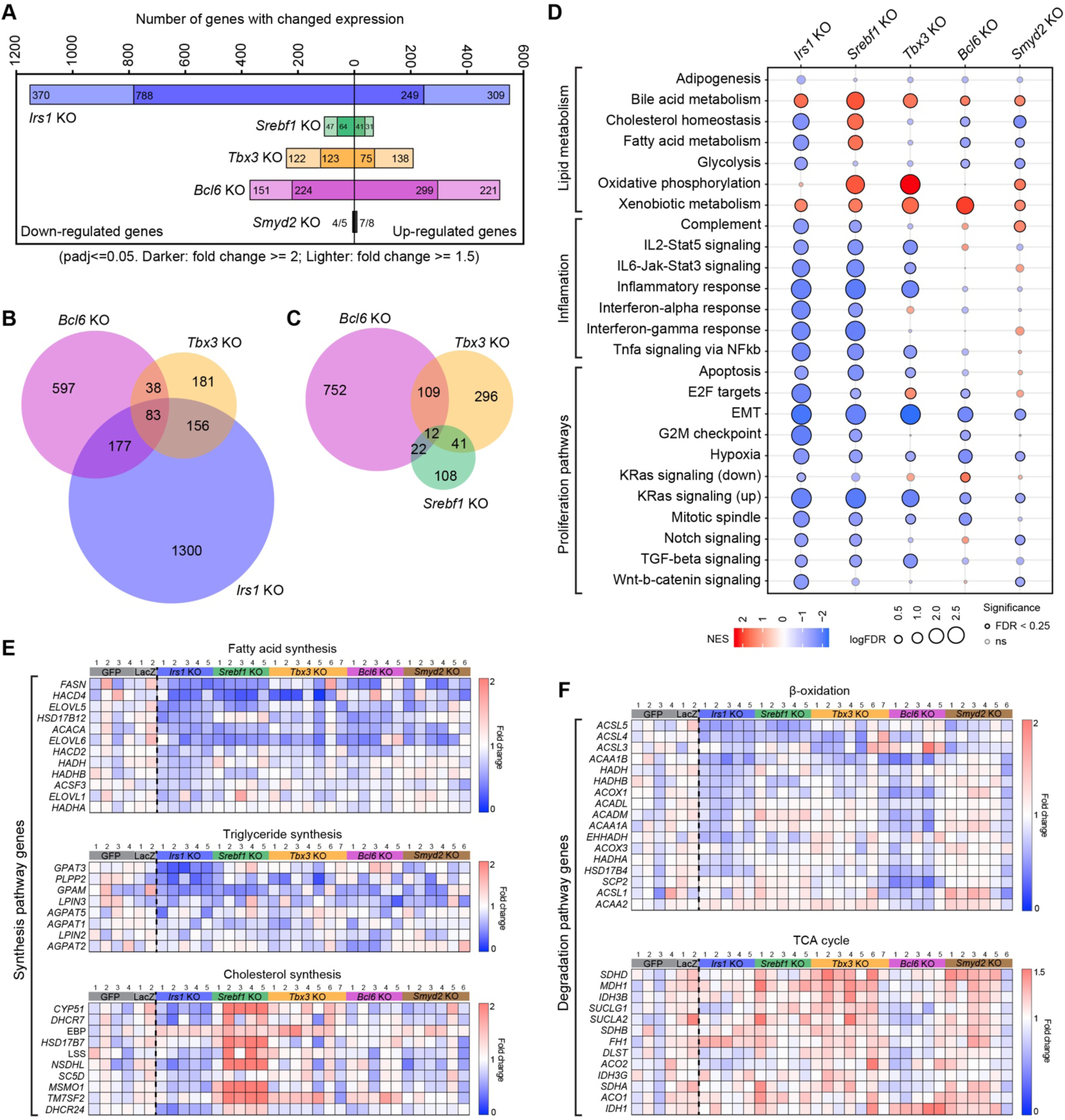
Transcriptional analysis of *Irs1*, *Srebf1*, *Bcl6, Tbx3*, and *Smyd2* KO livers after WD. A. The number of genes with altered expression in the RNA-seq data when comparing control (sgGFP and sgLacZ) and KO livers after 3 months of WD. Darker and lighter colored bars represent the number of differentially expressed genes with a fold change of >=2 and >=1.5, respectively. Genes with statistically significant fold change differences of less than 1.5 are not included here. B. Venn diagram showing the shared and unique gene numbers with changed expression (fold change >=1.5) in *Bcl6*, *Tbx3*, and *Irs1* KO livers. C. Venn diagram showing the shared and unique gene numbers with changed expression (fold change >=1.5) in *Bcl6*, *Tbx3*, and *Srebf1* KO livers. D. Hallmark pathway enrichment analysis of RNA-seq data from **A**. E. Heatmaps showing the fold changes of differentially expressed genes in fatty acid, triglyceride, and cholesterol synthesis pathways. The average expression level of control samples (four sgGFP and two sgLacZ) were normalized to 1 for each gene. F. Heatmaps showing the fold changes of differentially expressed genes in β-oxidation and TCA cycle pathways. The normalization method is the same as shown in **E**.

### A selective small molecule inhibitor of SMYD2 can prevent fatty liver disease

We sought to determine if chemical SMYD2 inhibition could serve as orthogonal validation of the genetic findings, and to investigate the therapeutic potential of SMYD2 blockade in fatty liver disease. We used AZ505, a selective SMYD2 inhibitor with an IC50 of 0.12 μM (Ferguson et al., 2011) that has strong selectivity for SMYD2 (>600 fold selectivity). WD was fed to WT mice at 8 weeks of age, followed by treatment with vehicle control or AZ505 with a dose of 10mg/kg (intraperitoneal injection) three times per week (Figure 7A and **Table S1**). This dose was well tolerated in our experiment. Two months after continued WD and inhibitor treatment, we observed significant reductions in body weight, liver weight, and liver/body weight ratios in AZ505 vs. vehicle treated mice (**Figure 7B-D**). Smaller, less pale livers were observed in AZ505 treated mice, and liver histology showed dramatically decreased steatosis in contrast to a wide range of macro and microscopic lipid droplet deposition in vehicle treated livers (Figure 7E). Accordingly, AZ505 treated mice showed improved transaminitis (**Figure 7F,G**) and a significant decrease in liver triglycerides (Figure 7H), whereas the liver cholesterol (Figure 7I) and plasma lipids (**Figure 7J,K**) showed a modest increase. Thus, a small molecule SMYD2 inhibitor can recapitulate the phenotypes of SMYD2 CRISPR KO mice. Because AZ505 binds to the active center of SMYD2 (Ferguson et al., 2011), these experiments also show that mechanistically, SMYD2 enzymatic activity, and not just protein levels, contributes to fatty liver phenotypes. In summary, several lines of orthogonal evidence suggest that SMYD2 is a promising NASH target.

**Figure 7.**
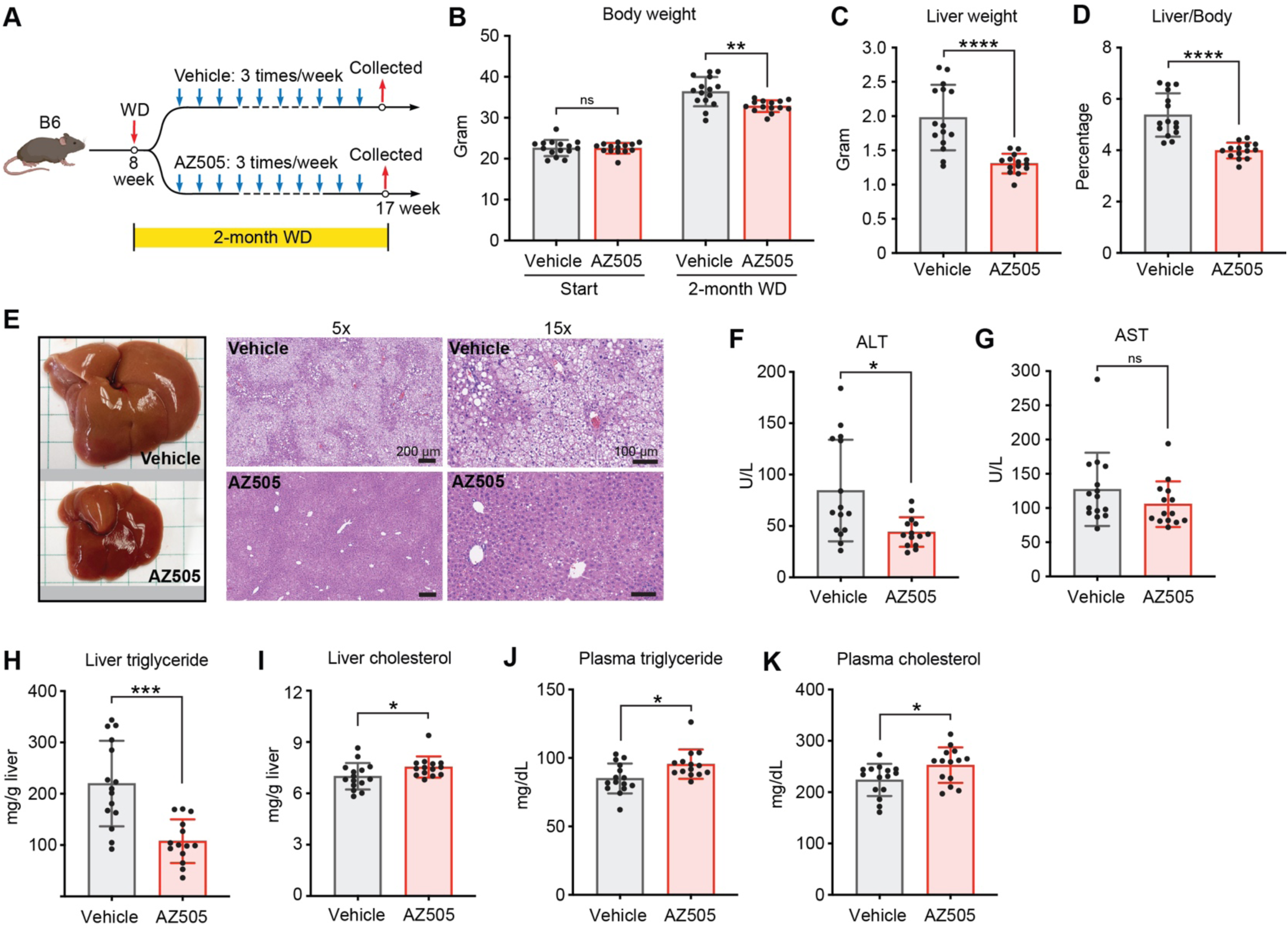
AZ505, a selective SMYD2 inhibitor, ameliorates fatty liver disease. A. Experimental design for pre-clinical testing of AZ505 in the WD NASH model. WD was given at 8 weeks of age for a total of 2 months. AZ505 treatment started one day after WD was initiated. Vehicle (2.5% DMSO in saline) or AZ505 (10 mg/kg in saline containing 2.5% DMSO) was given to mice intraperitoneally 3 times per week. B. Body weight of vehicle or AZ505 treated mice before and after 2 months of WD (n = 15 and 14 mice). C-D. Liver weight and liver/body weight ratios of vehicle or AZ505 treated mice after 2 months of WD (n = 15 and 14). E. Representative liver pictures and H&E staining of liver sections for vehicle or AZ505 treated mice after 2 months of WD. F-G. Liver function analysis using plasma ALT and AST (n = 15 and 14). H-I. Liver triglyceride and cholesterol analysis (n = 15 and 14). J-K. Plasma triglyceride and cholesterol analysis (n = 15 and 14).

## Discussion

The detection of mutant clone expansion and recurrent mutations within and between individuals provide compelling evidence for increased mutant clone fitness in chronic liver disease (Ng et al., 2021). Fatty liver disease has not been conceptualized as a disease of genetic mosaicism, so the biological impact of clonal heterogeneity has not yet been functionally explored. To explore this question for many genes at one time, we established the MOSAICS platform to model widespread somatic mosaicism in the absence of regeneration. The advance associated with this new technology is that it allows for *in vivo* screening with a much greater clone density, and in a non-proliferative setting, in contrast to our previous *Fah* KO screening model. Tracing of *Mboat7* and *Gpam* mutant hepatocytes and the results from the MOSAICS screens underscored the emerging concept that lipid laden cells have a fitness disadvantage when compared to lean cells. Thus, cells that acquire somatic mutations that lead to reduced lipid accumulation often become more fit than their WT counterparts. This new environmental pressure to reduce lipid accumulation fosters the selection of adaptive mutations within NASH livers.

Just because expanding mutant clones are more fit does not mean that the causative mutations, if applied to the entire tissue, can also increase organismal health. Because fitness comes in many forms, mutations can drive clonal expansion through diverse mechanisms that include increased proliferation, increased survival, or metabolic adaptations. Therefore, genomic data and *in vivo* screening in isolation were not sufficient to clarify how fitness is promoted, and whether or not this type of fitness is selfish or adaptive. It was important to perform tissue wide phenotypic analysis so that the impact of a mutation on the organismal level could be understood. We exploited a rapid whole liver KO method to accelerate genetic validation. This allowed multiple head to head comparisons between more than 20 different types of KO mice. We found that a significant subset of positively selected somatic mutations, when induced in the entire liver, led to improvements in fatty liver phenotypes. Interestingly, we showed that somatic mutations can be beneficial in the absence of increased proliferation or regeneration. It is likely that the clonal expansions associated with these mutations are caused by increases in cell survival, in addition to modest proliferation changes. In humans, these fitness improvements are enough to drive measurable levels of positive selection over decades. Remarkably, several lines of evidence suggest that these mutations do not promote cancer since many of these genes were not identified in large scale cancer sequencing efforts, and our *in vivo* experiments did not reveal greater rates of tumorigenesis.

Convergent and recurrent somatic mutations suggest that in disease conditions, humans are in effect performing pooled genetic screens that select for mutations that promote cell fitness in fatty liver disease. Several of the genetic hits are leading to adaptive mechanisms and potential therapeutic targets. This led us to hypothesize that sampling from a larger genetic space could uncover unexpected pathways in metabolic liver disease. Using evolutionary selection between genetically heterogeneous clones in cell culture systems has been a mainstay of cancer gene discovery. However, metabolic disease has not identified a facile method of gene discovery using phenotypic screens, *in vitro* or *in vivo*. Ultimately, it is hard to phenotype many genetic strains of mice or humans, but it is much easier to phenotype thousands of genetically distinct and competing cells. The MOSAICS platform identified multiple genes that when mutated, promote clonal fitness through the reduction of lipotoxicity. *Tbx3*, *Bcl6*, and *Smyd2* each have a significant impact on fatty liver development, and *Smyd2* inhibitors represent a promising approach for NASH treatment. These efforts validate the power of using high throughput somatic mutation selection as a gene discovery engine in metabolic disease.

Some of the most important human metabolic disease genes (*PCSK9, PNPLA3, HSD17B13*) have been identified using germline genetic methods such as GWAS and whole exome sequencing (Abul-Husn et al., 2018; Cohen et al., 2006; Romeo et al., 2008). However, germline mutations are inherently limited in the amount of genetic space that can be explored because lethal developmental phenotypes select against mutations that could have a benefit in specific tissues later in life. This study, in tandem with somatic sequencing studies from multiple labs, shows that somatic mutations are an alternate source for important disease genes. The positive selection of mutant clones in MOSAICS mice, and in chronically damaged human tissues, are new ways of identifying metabolic disease regulators and therapeutic targets. This could change the way we think about somatic mosaicism, becoming a source for genetic adaptation, rather than just a source of disease driving mutations.

## Acknowledgements

We thank Sam Wang and Andrew Hsieh for constructive comments on the manuscript; J. Shelton (UTSW Histopathology Core), E. Nwoka, and C. Moxon (UT Southwestern Tissue Management Shared Resource) for histopathology; D. Ramirez (UTSW Whole Brain Microscopy Facility) for whole liver imaging; J. Xu and Y. J. Kim (CRI Sequencing Core) for sequencing; S. Hacker, A. Walker, and J.I. Gamayot for metabolic phenotyping assays. T.W. is supported by NIH (R01CA258584). C.L. and the UT Southwestern Tissue Management Shared Resource are supported by the Simmons Comprehensive Cancer Center Support Grant (P30CA142543). Y.H. is supported by NIH (R01CA233794), CPRIT (RR180016), and the European Commission (ERC-AdG-2020-101021417, ERC-AdG-2014-671231). H.Z. is supported by the Pollack Foundation, the NIH (R01AA028791, R01DK125396), a Simmons Comprehensive Cancer Center Cancer & Obesity Translational Pilot Award, and the Emerging Leader Award from the Mark Foundation For Cancer Research (#21-003-ELA).

## Author contributions

Z.W. and H.Z. conceived the project, performed the experiments, and wrote the manuscript.

S.Z., N.F., and Y.H. performed bioinformatic analysis of human cirrhosis transcriptomic data and helped to conceptualize the paper.

N.K. reviewed the human NASH and cirrhosis liver histology.

R.G. performed the plasma assays and liver tissue lipid analysis.

C.L. oversaw tissue sectioning and histologic assays.

Y.W., Y.J., and T.W. performed bioinformatic analysis for mouse experiments.

M.Z. and T.S. generated *Gpam* floxed mice.

L.L., Q.Z., Y.L., and S.H. performed animal and cell culture experiments.

P.G. performed the histologic analysis and NASH scoring.

M.H. and P.C. provided conceptual contributions to the paper.

## Conflicts statement

H.Z. has a sponsored research agreement with Alnylam Pharmaceuticals, consults for Flagship Pioneering and Chroma Medicines, and serves on the SAB of Ubiquitix. H.Z., Z.W., and L.L. have a provisional patent on *GPAM* and *TBX3* siRNAs.

## METHODS

### Mouse strains and breeding

All mice were handled in accordance with the guidelines of the Institutional Animal Care and Use Committee at UT Southwestern. All experiments were done in an age and sex controlled fashion unless otherwise noted. C57BL/6 strain background mice were used for all the experiments. *Mboat7^tm1a(KOMP)Wtsi^/H* mice, obtained from KOMP, were originally generated by the Wellcome Trust Sanger Institute. *Gpam^f/f^* mice were generated in the CRI Mouse Genome Engineering Core at UT Southwestern. In brief, CAS9 protein, synthetic sgRNA, and single-stranded DNA containing *Gpam* exon3 flanked by LoxP sites and homology arms, were co-injected into C57BL/6 mouse zygotes, which were then implanted into CD-1 mice. Genotyping and Sanger sequencing was used to confirm homologous recombination in the genome-edited pups. *LSL-tdTomato* (strain #007914) and *Rosa-rtTA; TetO-Cas9* mice (#029415) were obtained from The Jackson Laboratory. Mice homozygous for both *Rosa-rtTA* and *TetO-Cas9* were used to ensure a high Cas9 expression level in the liver. Western Diet (WD) used for NASH modeling is described in (Tsuchida et al., 2018). It is composed of high fat solid food (ENVIGO #TD.120528) and high sugar water containing 23.1g/L d-fructose (Sigma-Aldrich #F0127) and 18.9 g/L d-glucose (Sigma-Aldrich #G8270).

### Lineage tracing in floxed mice

For whole liver deletion of *Mboat7* or *Gpam* gene, 5*10^10 genomic copies of commercially produced AAV8-TBG-Cre (Addgene #107787) or control AAV8-TBG-GFP (Addgene #105535) in 100μl saline was injected retro-orbitally into *Mboat7^f/f^* or *Gpam^f/f^* mice at 8 weeks of age. For mosaic deletion of *Mboat7* or *Gpam* and Tomato labeling of hepatocytes, 0.125*10^10 genomic copies of AAV8-TBG-Cre in 100μl saline was injected retro-orbitally into *Mboat7^f/f^; LSL-tdTomato* het, *Gpam^f/f^; LSL-tdTomato* het mice, or control *LSL-tdTomato* het mice at 8 weeks of age. One week after injection of low dose AAV8-TBG-Cre, 7 mice from each group were collected and analyzed for the initial labeling percentage in the liver. The remaining mice were divided into chow or WD groups and traced for another 4 or 6 months.

### Fluorescent imaging and image processing

For fluorescent imaging, liver pieces were fixed in buffered formalin (Fisherbrand #245-685) for 24h with gentle shaking at 4°C and then transferred into 30% sucrose (w/v) solution for another 24h with shaking at 4°C. The livers were then embedded and frozen in Cryo-Gel (Leica #39475237), and sectioned at a thickness of 16μm. Images were taken using a Zeiss Axionscan Z1 system in the UTSW Whole Brain Microscopy Facility to visualize and quantify Tomato clones. To statistically analyze the percentage of Tomato positive cells, black and white fluorescent images were taken from the same slide using an Olympus IX83 microscope at 4x magnification. Two different fields were taken for each liver. The percentage of Tomato positive cells (bright areas) was analyzed using ImageJ.

### H&E, immunohistochemistry (IHC) and TUNEL staining

Liver pieces were fixed in buffered formalin (Fisherbrand #245-685) for 24h with gentle shaking at 4°C and then transferred to 70% EtOH for another 24h with shaking at 4°C. Paraffin embedding, liver sectioning (4μm thickness), and H&E staining were performed at the UT Southwestern Tissue Management Shared Resource Core. For IHC staining, the following primary antibodies were used: Pten (CST #9559, IHC 1:200); Ki67 (Abcam #AB15580, IHC 1:200). IHC was performed as previously described (Zhu et al., 2010). TUNEL staining was performed on paraffin embedded liver sections using *In Situ* Cell Death Detection Kit, Fluorescein (Roche #C755B40) according to the manufacturer’s protocol. QuPath software (https://qupath.github.io/) was used to quantify Ki67 and TUNEL staining.

### Plasma parameters and liver lipid measurements

Blood samples were taken using heparinized tubes from the inferior vena cava immediately after sacrificing the mouse, and then transferred into 1.5ml tubes and centrifuged at 2000g for 15min at 4°C. The supernatant after centrifugation (plasma) was analyzed for AST, ALT, cholesterol, and triglyceride using VITROS MicroSlide Technology at the UT Southwestern Metabolic Phenotyping Core. 100-150mg of liver per sample was weighted and used for lipid extraction and quantification at the UT Southwestern Metabolic Phenotyping Core.

### MOSAICS reagent construction

The MOSAICS plasmid uses the pX602 plasmid as a backbone. The sequence between the two AAV ITRs were removed using the NsiI and NotI restriction enzymes. The following fragments were cloned between the two AAV ITRs: the first SB100 binding IR, a U6 driven sgRNA scaffold, a CAG promoter driven SB100-P2A-Cre fusion cDNA with a beta-globin poly(A) signal, and the second SB100 binding IR. For library construction, mouse candidate genes for all of the *in vivo* screens were generated by using the mouse homologs of the human genes shown in **Table S3**. A few genes were not included in the mouse gene lists due to the lack of a homolog or because they were known tumor suppressor genes. The individual sgRNA sequences corresponding to mouse candidate genes were extracted from the Brie library (Doench et al., 2016) or obtained from the GUIDES server (http://guides.sanjanalab.org/), and synthesized by CustomArray. Most genes have 5 distinct sgRNAs, 4 from Brie and 1 from GUIDES. A few genes have 4 targeting sgRNAs due to the overlap of sgRNA sequences from Brie and GUIDES. See **Table S2** for the sgRNA sequences of targeted genes. The library construction protocol was described in (Canver et al., 2018). Each sgRNA maintained a >1000-fold representation during construction. For individual sgRNA cloning, forward and reverse primers were annealed and fused to BsaI digested MOSAICS plasmid using T4 ligase. See **Table S5** for the primers associated with individual sgRNAs used in this paper.

### AAV production and purification

AAV8 was produced using AAV-Pro 293T cells (Takara #632273) cultured in one or more 15cm dishes. Cells were plated one day before transfection at 50% confluence, which would allow the cells to reach 80-90% confluence the next day. For transfection of one 15cm dish, 10μg MOSAICS vector, 10μg pAAV2/8 (Addgene #112864) and 20μg pAdDeltaF6 (Addgene #112867) plasmids were mixed with 1ml Opti-MEM medium in one tube. In another other tube, 160μl PEI solution (1mg/ml in water, pH7.0, powder from ChemCruz #sc-360988) was mixed with 1ml Opti-MEM medium. The solutions from both tubes were then mixed and put aside for 10min before adding to cell culture. 48h after transfection, the cells were scraped off the dish and collected by centrifugation at 500g for 10min. The supernatant was disinfected and discarded, and the cell pellets were lysed in 2ml/15cm dish lysis buffer (PBS supplemented with NaCl powder to final concentration of 200mM, and with CHAPS powder to final concentration of 0.5% (w/v)). The cell suspension was put on ice for 10min with intermittent vortexing, and then centrifuged at 20,000g for 10min at 4°C. The supernatant containing the AAV was collected. To set up the gravity column for AAV purification, 0.5ml of AAV8-binding slurry beads (ThermoFisher #A30789), enough to purify AAV from three 15cm dishes, was loaded into an empty column (Bio-Rad #731-1550). After the beads were tightly packed at the bottom, they were washed with 5ml of wash buffer (PBS supplemented with NaCl powder to a final concentration of 500mM). The supernatant containing AAV was then loaded onto the column. After all of the supernatant flowed through, the beads were washed with 10ml wash buffer twice. The AAV was then eluted with 3ml elution buffer (100mM glycine, 500mM NaCl in water, pH 2.5) and the eluate was immediately neutralized with 0.12ml 1M Tris-HCl (pH 7.5-8.0). The AAV was concentrated by centrifugation at 2000g for 3-5min at 4°C using an 100k Amicon Ultra Centrifugal Filter Unit (Millipore #UFC810024). After centrifugation, the volume of AAV should be equal to or less than 0.5ml. The concentrated AAV was diluted with 4-5ml AAV dialysis buffer (PBS supplemented with powders to final concentrations of 212mM NaCl and 5% sorbitol (w/v)) and centrifuged at 2000g for 3-5min at 4°C. The dilution and centrifugation processes were repeated 3 times. The final concentrated AAV was transferred into a 1.5ml tube and centrifuged at 20,000g for 5min to remove debris. The supernatant was aliquoted, flash frozen using liquid nitrogen, and stored at -80°C.

### MOSAICS screening and single gene deletion in inducible Cas9 mice

To functionally validate the MOSAICS vector, a *Pten* sgRNA (see **Table S5** for the *Pten* sgRNA primers) was cloned into the vector and the corresponding AAV8 was produced. Different volumes of concentrated AAV8-*sgPten* were diluted to a final volume of 100μl using saline, then retro-orbitally injected into *Rosa-rtTA; TetO-Cas9* double homozygous mice 3 days after dox water (1mg/ml dox) was initiated at 6.5 weeks of age. After determination of *Pten* KO efficiency using IHC and confirmation of SB100-Cre fusion protein expression and recombination using fluorescence imaging with *LSL-tdTomat*o mice (Figure 2C), the concentration of this AAV8-*sgPten* was used as a standard. The relative concentrations of the other AAVs, including library AAVs and single gene targeting AAVs were all compared to AAV8-*sgPten* by qPCR using a pair of primers within the Cre sequence of the vector (See **Table S5** for primer sequences). The absolute genomic copy numbers were not determined. For the AAV libraries used for NASH, transcription factor, and epigenetic factor screens, a volume corresponding to 7μl of AAV8-*sgPten* (**Figure 2D,E**) was injected. For the AAV mini pool containing 8 sgRNAs (Figure 3E), a volume corresponding to 1μl of AAV8-*sgPten* (**Figure 2D,E**) was injected. For AAVs targeting individual genes (**Figure S6A**), a volume corresponding to 20μl AAV8-*sgPten* (**Figure 2D,E**) was injected. All AAVs were diluted to a final volume of 100μl using saline prior to injection. Dox water was withdrawn 10 days after injection and then chow or WD was given to mice for the specified times.

### Genomic DNA extraction, sgRNA amplification, and amplicon library construction

To extract genomic DNA containing the integrated sgRNA, the whole liver (except a small piece used for sectioning and H&E staining) was minced into about 1mm^3^ pieces using a blade and weighed. Small nodules observed in some epigenetic factor screening livers given WD (<=3 nodules per liver in 5 out of 8 livers) were excluded from samples being processed for genomic DNA extraction. Minced[3] liver in two volumes (w/v) of homogenizing buffer (100mM NaCl, 25mM EDTA, 0.5% SDS, 10mM Tris-HCl, pH 8) was transferred into a glass Wheaton Dounce Tissue Grinder[4] and stroked for 50 times or until no bulk tissues were seen. After homogenizing, 200μl chow fed liver lysate or 300μl WD-fed liver lysate was transferred to a 15ml tube for genomic DNA extraction using the Blood & Cell Culture DNA Midi Kit (Qiagen #13343) according to the manufacturer’s protocol. The remaining lysates were frozen in -80°C as backup samples. Briefly, 10ml Buffer G2 from the kit, 100μl Proteinase K (Roche #03115828001, or Proteinase K from the Qiagen kit) and 100μl RNase A (Invitrogen #12091-021) were added to the 15ml tube containing the lysate and digested in a 50°C water bath overnight. The next day, the tubes were centrifuged at 4000g for 10min and the lipid layer on the top was discarded. The remaining supernatant was loaded on the column, washed, and genomic DNA elution/precipitation were performed according to the manufacturer’s protocol. The precipitated DNA was resuspended in 100μl 10mM Tris (pH 8.0) and shaken on a 55°C shaker for 2h to help it dissolve. The amplicon library preparation protocol for high-throughput sequencing was adapted from (Canver et al., 2018). Briefly, 5μg genomic DNA, 5μl general forward primer mix (5μM), 5μl barcode specific reverse primer (5μM), 1μl Q5 DNA polymerase, 10μl Q5 buffer, 10μl HighGC buffer, 1μl dNTP, and water was mixed for a 50μl PCR reaction, and two reactions were made for each genomic DNA sample. The PCR cycle was 95°C 3min-(95°C 30s-60°C 30s-72°C 20s)*n-72°C 2min. The PCR cycle number was pre-optimized using the same PCR reactions with a smaller volume. The cycle numbers that gave a weak but sharp band on the DNA gel were used. In the final PCR reaction, 23 cycles were used for preparing the NASH gene, transcription factor, and epigenetic factor screens, and 30 cycles were used for preparing the 8 guide mini pool validation screen. After PCR, the two tubes of reactions with the same genomic DNA template were combined (total 100μl) and 70μl was resolved on a DNA gel. The 250bp band corresponding to the amplicon was cut and purified using the QIAquick Gel Extraction Kit (Qiagen #28704). The DNA concentration was determined using Qubit kit (Invitrogen #Q32853) and high-throughput sequencing was performed using an Illumina NextSeq500 system at the CRI at UT Southwestern Sequencing Facility.

### Bioinformatic analysis of MOSAICS screening results

The reads from the sequencing of amplicon libraries described above were trimmed with cutadapt (version 1.9.1) to remove the excessive adaptor sequences so that only the sgRNA sequences were retained. The 5’ sequences were trimmed with the options -O 32 --discard-untrimmed -g CTTTATATATCTTGTGGAAAGGACGAAACACCG. The 3’ sequences were trimmed with the options -O 12 -a GTTTTAGAGCTAGAAATAGCA. The abundance of each sgRNA was calculated with count function in MAGeck (version 0.5.6) with the default options(Li et al., 2014). The enrichment of each sgRNA was calculated with the test function in MAGeck with default options.

### RNA-seq library preparation and transcriptome analysis of mouse fatty livers

Total liver RNA was extracted from 4 sgGFP, 2 sgLacZ, 5 *Irs1*, 5 *Srebf1*, 7 *Tbx3*, 5 *Bcl6* and 6 *Smyd2* KO livers using TRIzol reagent (Invitrogen #15596026) followed by purification using the RNeasy Mini kit (Qiagen #74014). Briefly, a liver fragment with a size of about 3*3*3 mm^3^ from each sample was homogenized in 1ml Trizol, followed by adding 200μl chloroform and vortexed. After centrifugation at 20,000g for 10min at 4°C, 350μl supernatant from each sample was transferred to a new tube and mixed with equal volume of 75% EtOH, and then loaded on an RNeasy column. The following wash steps using RW1 and RPE buffers and RNA precipitation step were performed according to the manufacturer’s protocol. RNA-seq libraries were prepared with the SMARTer Stranded Total RNA Sample Prep Kit - HI Mammalian (Takara #634875). 75 bp single-end sequencing was performed using an Illumina NextSeq500 system at the CRI at UT Southwestern Sequencing Facility. Alignment, quantification, and differential expression analysis were performed using the QBRC_BulkRnaSeqDE pipeline (https://github.com/QBRC/QBRC_BulkRnaSeqDE). Briefly, the alignment of reads to the mouse reference genome (mm10) was done using (v2.7.2b) (Dobin et al., 2013). FeatureCounts (v1.6.4)(Liao et al., 2014) was then used for gene count quantification. Differential expression analysis was performed using the R package DEseq2 (v1.26)(Love et al., 2014). Cutoff values of absolute fold change greater than 2 and FDR<0.05 were used to select for differentially expressed genes between sample group comparisons. Finally, GSEA was carried out with the R package fgsea (v1.14.0) using the ’KEGG’ and ’Hallmark’ libraries from MsigDB.

### Transcriptome analysis of human NASH livers

Dataset 1 (NASH cohort): We downloaded the RNA-Seq transcriptome profiles of biopsied liver tissues (GEO: GSE130970) (Hoang et al., 2019) and analyzed 72 NASH patients with a range of disease severities (NASH activity scores of 1 to 6) and relevant histological features, i.e., steatosis, inflammation, fibrosis, and hepatocyte ballooning. The raw sequence reads were aligned to the GENCODE human reference genome (GRCh37, p13) using the STAR aligner (ver 2.6.1b) (Dobin et al., 2013), and gene-level count data were generated by the featureCounts function in the Subread package (ver 1.6.1)(Liao et al., 2014) and the GENCODE genome annotation (GRCh37, v19) (Frankish et al., 2019). The count data were normalized using “Relative Log Expression” normalization (RLE) implemented in the DESeq2 package (Love et al., 2014). Dataset 2 (HCV cirrhosis cohort): We analyzed the microarray gene expression profile of formalin-fixed needle biopsy specimens from the livers of 216 patients with hepatitis C-related early-stage (Child-Pugh class A) cirrhosis (GEO: GSE156540) (Hoshida et al., 2013). This cohort was prospectively followed for a median of 10 years at an Italian center with relevant time event outcomes collected, including child, death, HCC and decomposition.

### Drug treatment

AZ505 powder was purchased from MCE (#HY-15226). 40mg/ml AZ505 in 100% DMSO stock solution was made, aliquoted, and stored at -80°C. Before use, the stock solution was diluted 40x using 0.9% sodium chloride (Baxter #2F7123). WD was given to B6 mice at 8 weeks of age for a total of 2 months. AZ505 or vehicle (0.9% sodium chloride containing 2.5% DMSO) treatment started one day after WD and was administered IP at a dose of 10mg/kg three times per week until one day before euthanasia and collection of livers.

### Curating transcription and epigenetic factors with putative pathogenic activity in NASH

Gene expression of transcription factors do not directly reflect their functional activities, due to the low correlation between gene expression and protein abundance as well as the co-regulation with co-factors. Transcription factor activities can be estimated using enrichment of their downstream targets (Liberzon et al., 2011; Subramanian et al., 2005). However, as we observed in our previous study (Topol et al., 2016), traditional methods such as Gene Set Enrichment Analysis (GSEA) overestimated the number of significant transcription factors, largely confounded by the large number of overlaps among putative target genes of different TFs (Friedman et al., 2009). Therefore, we performed global modeling of putative transcription factors (Balwierz et al., 2014) with gene expression data from NASH (GEO: GSE130970) (Hoang et al., 2019) and HCV cirrhosis patients (GEO: GSE15654) (Hoshida et al., 2013), to directly infer the transcription factor activities from their downstream targets with adjustment for overlapping targets. A similar linear regression-based model was previously proposed to predict transcription factor regulatory activities and motifs from yeast gene expression data (Conlon et al., 2003). The method regresses the fold-change of a gene on its putative regulatory transcription factor(s). The coefficient (Z score) of a transcription factor, estimated using genome-wide fold changes and predicted targets of all transcription factors, represents the regulatory activity change of the transcription factor across all the liver patients.

The regression model is defined as following:

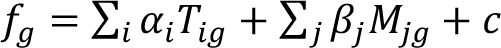

where *f_g_* is the fold change of *g-th* gene between two conditions; *T_ig_* is the number of binding sites of *i-th* TF on the promoter of the *g-th* gene; *M_ig_* is the number of binding sites of the *j-th* microRNA on the 3’ UTR of the *g-th* gene; and α_*i*_, β_*j*_ and *c* (a constant) can be inferred based on the values of *f_g_ T_ig_* and *M_jg_* for all the genes in the RNA-seq data. The Z scores of coefficients α_*i*_ and β_*j*_ represent the activity changes of the *i-th* TF and j-th microRNA. Global transcription factor binding sites represented by 190 position-weighted matrices (PWMs) covering 500 mammalian TFs were based on the union of JASPAR (Wasserman and Sandelin, 2004), TRANSFAC (Matys et al., 2003), and additional motifs from chromatin immunoprecipitation with DNA microarray and ChIP-seq data collected by Balwierz et al. (Balwierz et al., 2014). The initial regression analysis was done using ISMARA before further integrative analysis with patient clinical histological features and time event outcome; sample-specific transcription factor activity was estimated by the same regression model, where the fold changes were calculated between a single sample and all the samples combined together.

In the NASH cohort, the sample-level activities for each TF were associated with the four histological features, including fibrosis, inflammation, ballooning, and steatosis based on Pearson correlation. In the HCV cirrhosis cohort, the activities for each TF were used to perform outcome analysis on four time events, including child, death, HCC and decomposition using cox proportional regression model. The p-values were calculated for both analyses respectively, followed by the calculation of False Discovery Rate (FDR) for multiple testing correction.

Similarly, to screen the putative pathogenic epigenetic regulators, we also calculated the Pearson correlation between the gene expression of epigenetic regulators and the four histological features (fibrosis, inflammation, ballooning, and steatosis) in the NASH cohort, and used gene expression of known epigenetic regulators as independent variables to perform the cox proportional regression on four time events (Child-Pugh, death, HCC and decompensation) in the HCV cirrhosis cohort. The p-values were calculated for both analyses respectively, followed by the calculation of FDR for multiple testing correction.

## Statistical Analysis

The data in most panels reflect multiple experiments performed on different days using mice derived from different litters. Variation in all panels is indicated using standard deviation presented as mean ± SD. Two-tailed unpaired Student’s t-tests were used to test the significance of differences between two groups. Statistical significance is displayed as ns (not significant, or p>=0.05), * (p<0.05), ** (p<0.01), *** (p<0.001), **** (p<0.0001) unless specified otherwise. Image analysis for quantification was blinded.

## Materials availability

Plasmids, mouse lines, and other unique reagents generated in this study will be distributed upon request after completion of relevant material transfer agreements.

## Data and code availability

RNA-seq raw data will be deposited to GEO and made publicly available prior to publication. The analyzed results are in **Table S4**. Raw data for all the graphs in the figures are available in **Table S4**. There is no original code in this paper. Any additional information required to reanalyze the data in this paper will be made available from the lead contact upon request.

**Supplemental Table S1. Mouse phenotyping data.**

Sheet 1. *Mboat7* whole-liver KO and control mice fed with 1.5 months of chow or WD. Sheet 2. *Gpam* whole-liver KO and control mice fed with 3 months of chow or WD. Sheet 3. MOSAICS based single gene whole liver KO mice.

Sheet 4. Drug treated mice.

**Supplemental Table S2. MOSAICS screening data.**

Sheet 1. Results for 63 NASH gene screening (genes). Sheet 2. Results for 63 NASH gene screening (sgRNAs).

Sheet 3. Results for 8 guide mini pool screening, related to Figure 3F. Sheet 4. Results for transcription factor screening (genes).

Sheet 5. Results for transcription factor screening (sgRNAs). Sheet 6. Results for epigenetic factor screening (genes).

Sheet 7. Results for epigenetic factor screening (sgRNAs).

**Supplemental Table S3. Human transcription factor and epigenetic factor candidates.**

Sheet 1. Fatty liver disease associated transcription factors identified in NAFLD patients. Sheet 2. Fatty liver disease associated transcription factors identified in fibrotic HCV patients. Sheet 3. Fatty liver disease associated epigenetic factors identified in NAFLD patients.

Sheet 4. Fatty liver disease associated epigenetic factors identified in fibrotic HCV patients.

**Supplemental Table S4. RNA-seq data from single gene KO mouse models.**

Sheet 1. FPKMs from 4 sgGFP, 2 sgLacZ, 5 *Irs1*, 5 *Srebf1*, 7 *Tbx3*, 5 *Bcl6* and 6 *Smyd2* KO livers from MOSAICS based whole liver KO mice.

Sheet 2. Expression changes for *Irs1* KO livers vs. control livers (sgGFP and sgLacZ) Sheet 3. Expression changes for *Srebf1* KO vs. control livers (sgGFP and sgLacZ).

Sheet 4. Expression changes for *Tbx3* KO vs. control livers (sgGFP and sgLacZ). Sheet 5. Expression changes for *Bcl6* KO vs. control livers (sgGFP and sgLacZ). Sheet 6. Expression changes for *Smyd2* KO vs. control livers (sgGFP and sgLacZ).

**Supplemental Table S5. Primers used in this study.**

**Figure S1.**
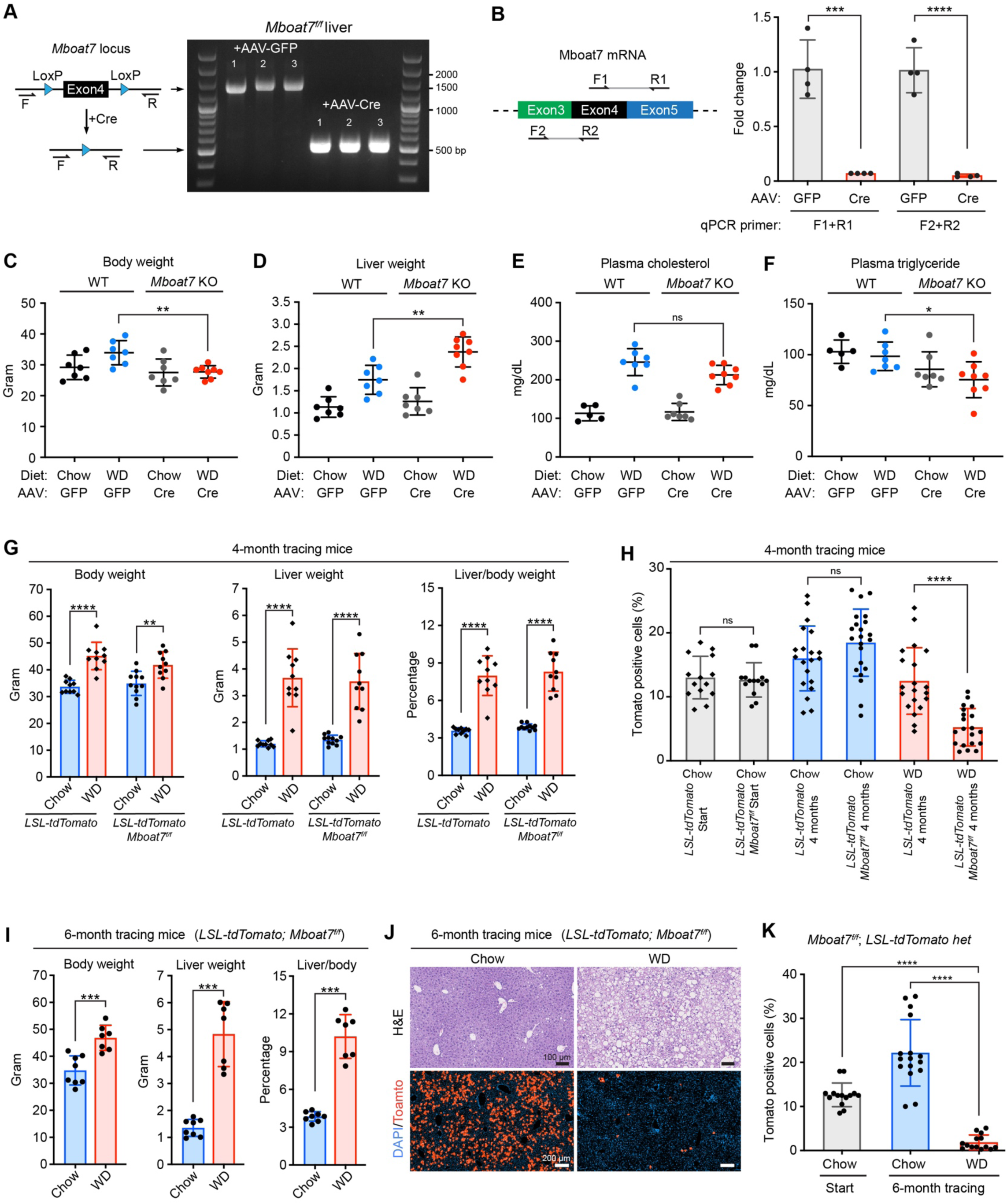
Characterization of whole-liver and mosaic *Mboat7* KO mice. A. Genotyping of *Mboat7^f/f^* liver tissues before and after Cre recombination (n=3 and 3). B. Examination of *Mboat7* mRNA levels in the liver before and after Cre recombination. Primer design is shown on the left. Two pairs of primers were used (n = 4 for each group). C-D. Body and liver weights of whole-liver *Mboat7* KO mice treated with chow or WD and their corresponding control mice (n = 7, 7, 7, 8 mice for each group). E-F. Plasma triglyceride and cholesterol analysis for the mice in **C** (n = 5, 7, 7, 8 for each group). G. Body weight, liver weight, and liver/body weight ratios of *LSL-tdTomato* het mice and *Mboat7^f/f^; LSL-tdTomato* het mice in a 4-month lineage tracing experiment as described in Figure 1G (n = 10, 10, 11, 10 mice for each group). H. Statistical comparison of the data from Figure 1I and **1J**. I. Body weight, liver weight, and liver/body weight ratios of *Mboat7^f/f^; LSL-tdTomato* het mice in a 6-month lineage tracing experiment. *Mboat7^f/f^; LSL-tdTomato* het mice were injected with low-dose of AAV8-TBG-Cre and treated with chow or WD for 6 months (n = 8 and 7 mice for each group). J. Representative H&E and fluorescent pictures of liver sections in the 6-month tracing experiment. K. Statistical analysis of Tomato-positive cells for the liver sections of *Mboat7^f/f^; LSL-tdTomato* het mice in **J** (n = 8 and 7 mice for each group). Each dot represents one image field; two different fields in each liver were analyzed. The same starting point Tomato labeling data as was included in Figure 1J are included here for the purpose of comparison.

**Figure S2.**
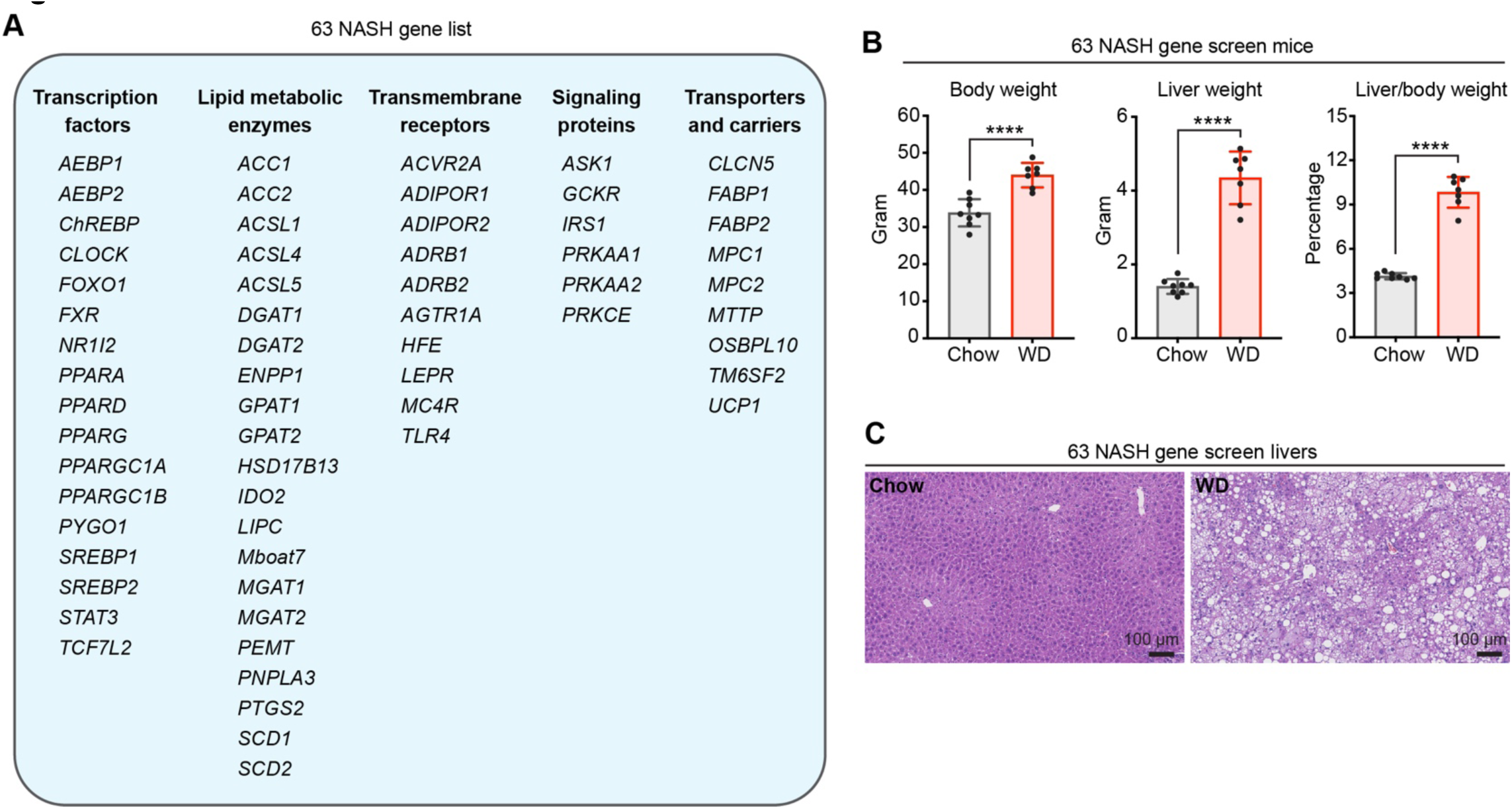
Tracing of mosaic mutant hepatocytes and validation of the most enriched genes. A. List of the targeted NASH genes in the MOSAICS experiment described in Figure 3A. B. Body weight, liver weight, and liver/body weight ratios of mice at the endpoint of the MOSAICS experiment described in Figure 3A (n = 8 and 7 mice for each group). C. Representative H&E staining of liver sections for the mice in **B**.

**Figure S3.**
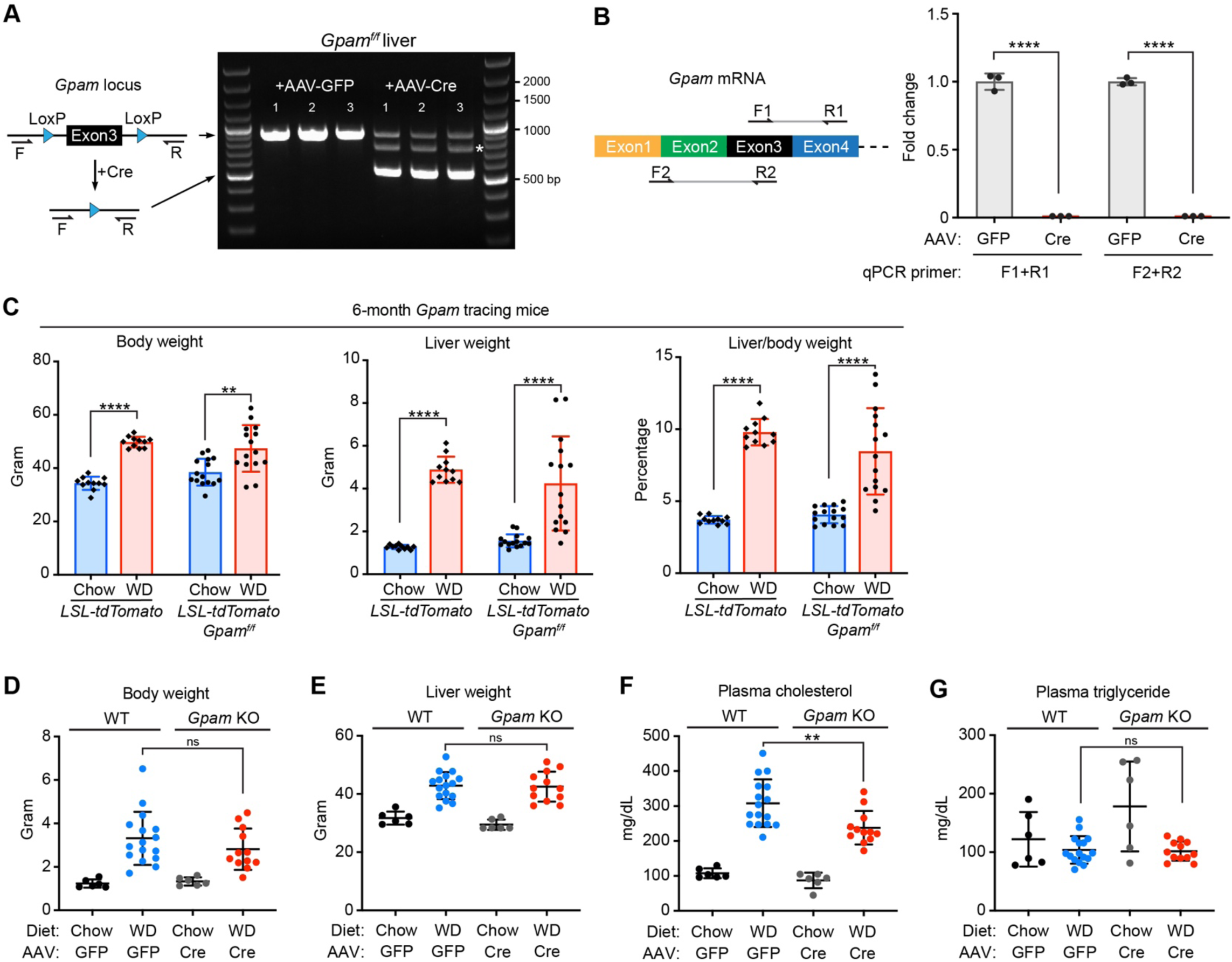
Characterization of whole-liver and mosaic *Gpam* KO mice. A. Genotyping of *Gpam^f/f^* liver tissues before and after Cre recombination. The asterisk is next to a hybrid annealing band from the top and bottom PCR products. B. qPCR examination of *Gpam* mRNA levels in livers before and after Cre recombination. Primer design is shown on the left. Two pairs of primers were used (n = 3 mice for each group). C. Body weight, liver weight, and liver/body weight ratios of *LSL-tdTomato* het mice and *Gpam^f/f^; LSL-tdTomato* het mice in the 6-month lineage tracing experiment as described in Figure 4A. D-E. Body and liver weights of whole-liver *Gpam* KO mice treated with chow or WD and their corresponding control mice (n = 6, 16, 6, 12 mice for each group). F-G. Plasma cholesterol and triglyceride analysis for the mice in **D-E** (n = 6, 16, 6, 12 for each group).

**Figure S4.**
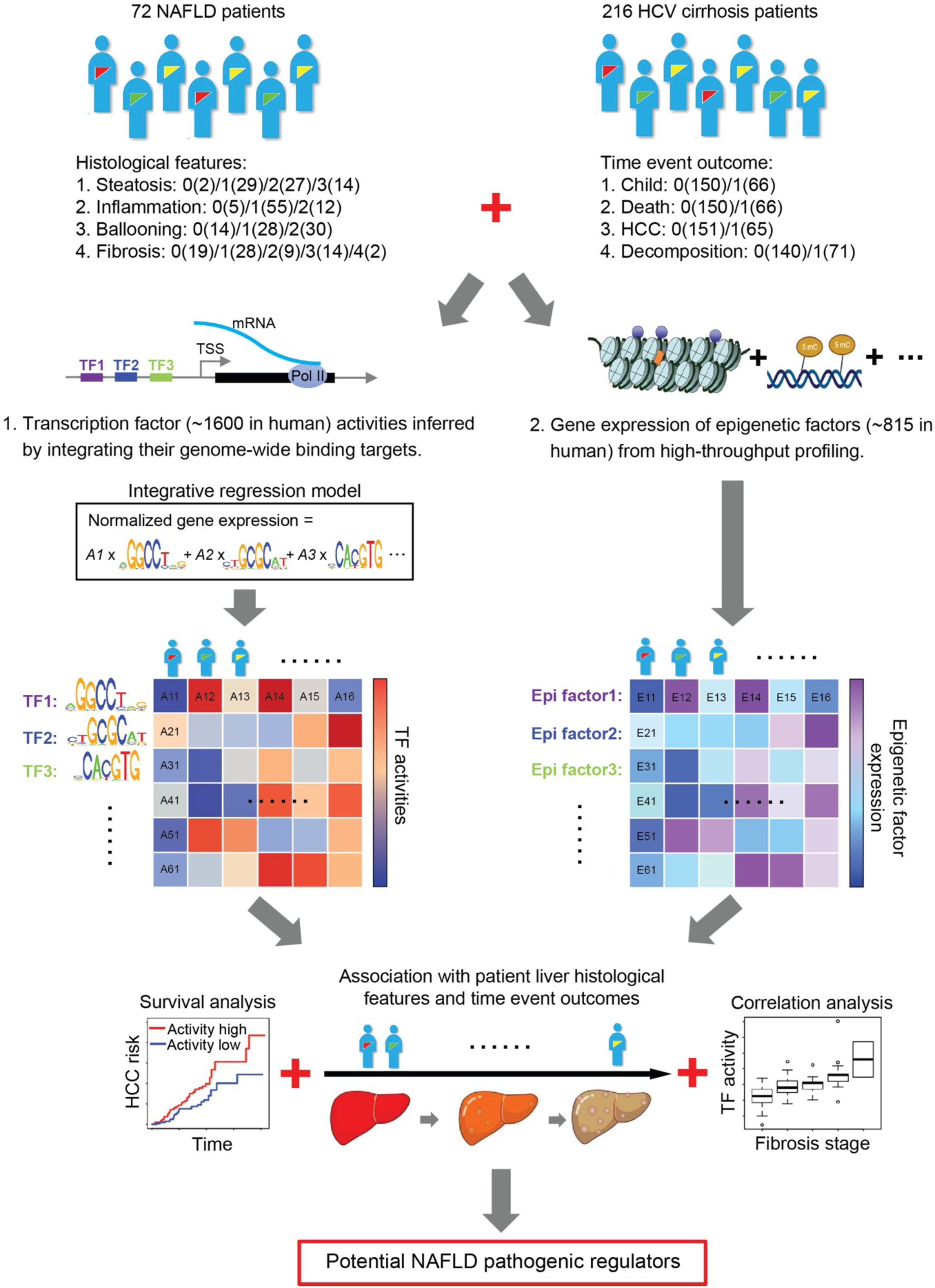
Workflow for identifying differentially activated (for transcription factors) or expressed (for epigenetic proteins) proteins in NAFLD and HCV cirrhosis patients.

**Figure S5.**
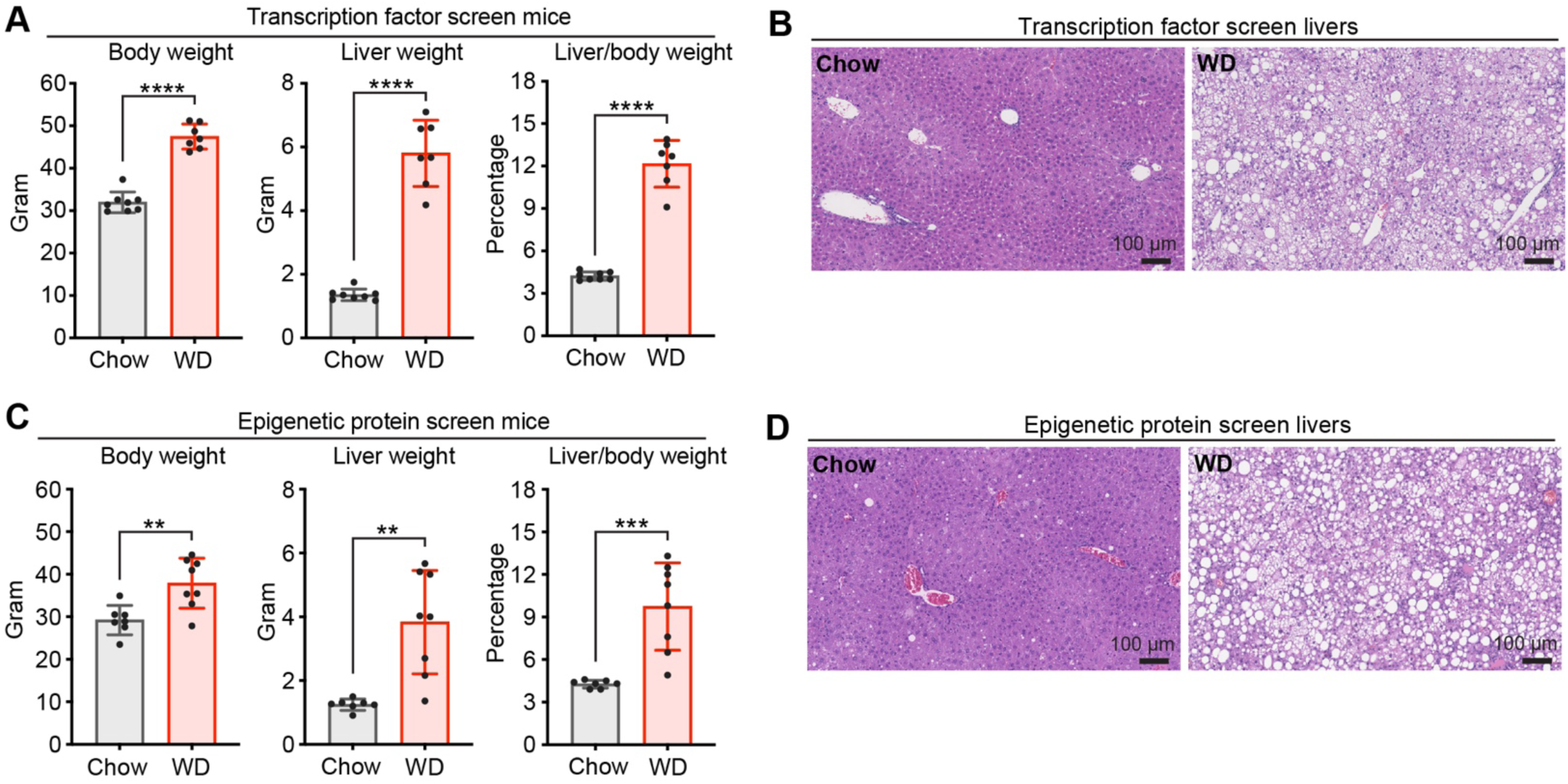
Characterization of MOSAICS livers after 6 months of screening. A. Body weight, liver weight, and liver/body weight ratios of mice at the endpoint of the transcription factor screen (n = 8 and 7 mice for each group). B. Representative H&E staining of liver sections for the mice in **A**. C. Body weight, liver weight, and liver/body weight ratios of mice at the endpoint of the epigenetic factor screen (n = 7 and 8 mice for each group). D. Representative H&E staining of liver sections for the mice in **C**.

**Figure S6.**
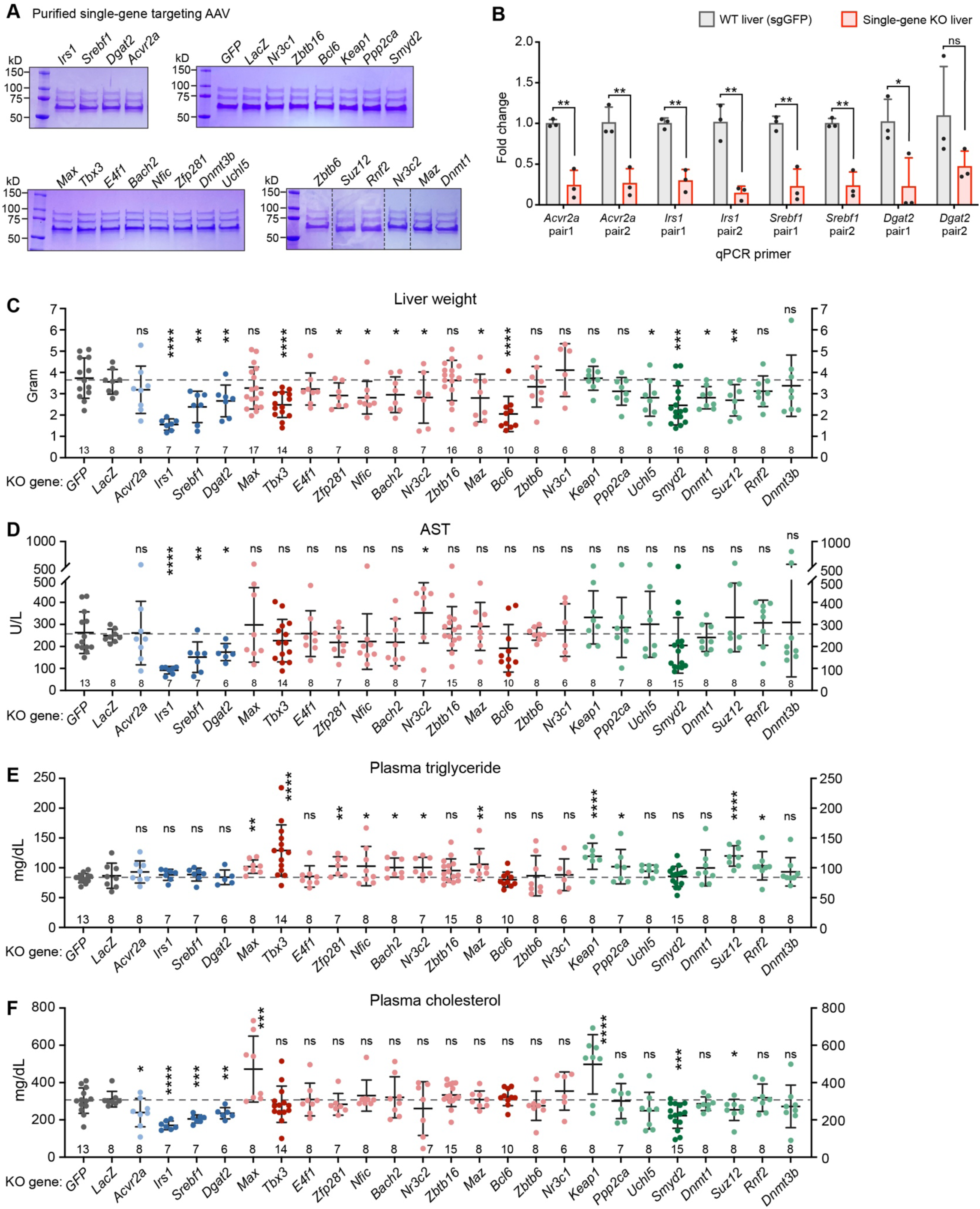
Characterization of liver-specific KO phenotypes for the top hits from somatic mosaic screening. A. Purified MOSAICS AAVs carrying one sgRNA used for generating single-gene whole liver KO mice. B. qRT-PCR assays to measure target gene KO efficiencies. In each primer pair, either forward or reverse primer spans the Cas9 cutting site of the indicated gene, and thus cannot anneal to the edited cDNA reverse transcribed from whole liver mRNA collected after 10 days of Dox water. The red column represents the ratio of intact mRNA of the indicated gene in the KO vs. WT livers (n=3 for each group). C. Liver weight of control (sgGFP and sgLacZ) and liver-specific KO mice fed with 3 months of WD. The color scheme is the same as in Figure 5E-I. Each dot represents one mouse. The n is denoted at the bottom of each plot. D-F. Liver function analysis using plasma AST, plasma triglyceride, and plasma cholesterol.

**Figure S7.**
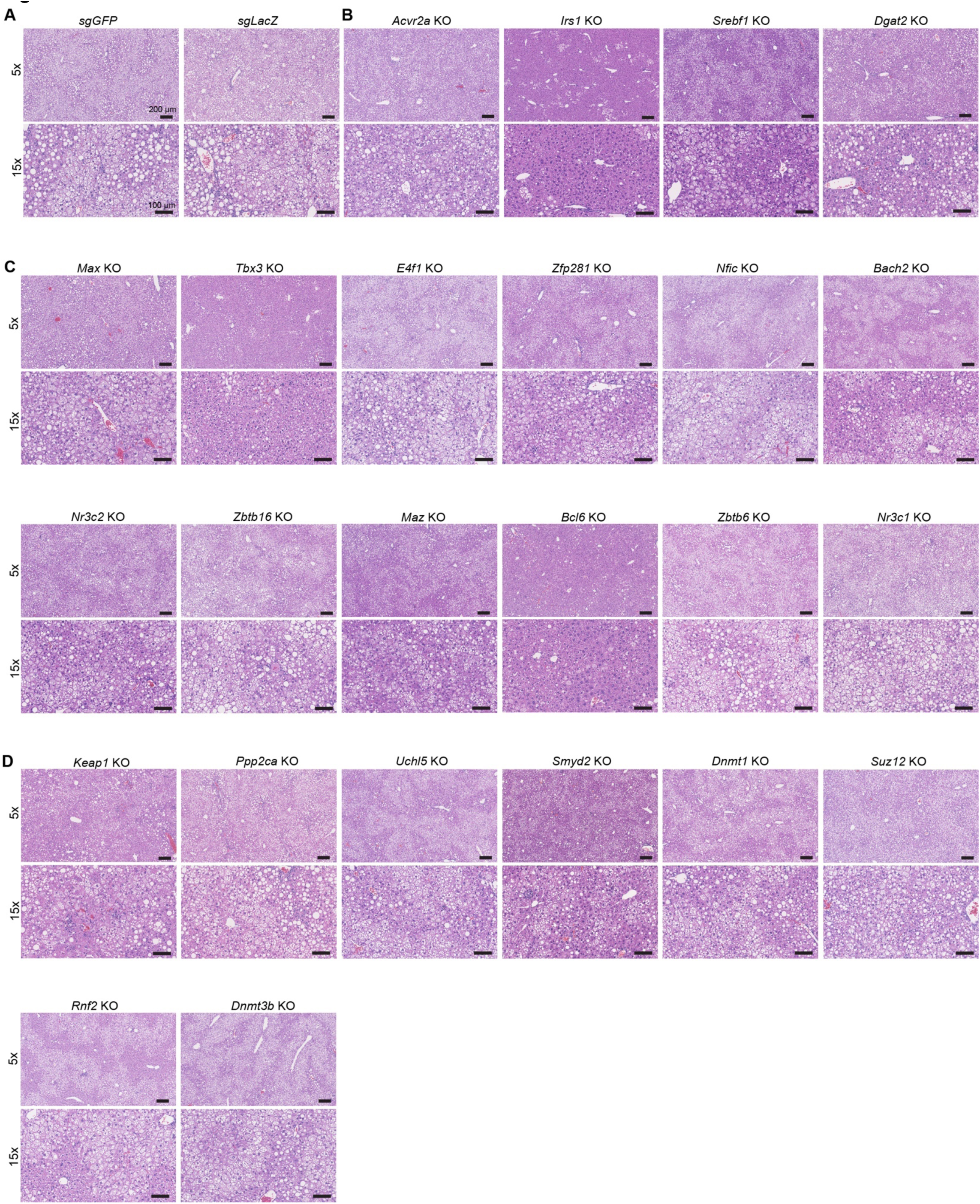
Representative H&E staining of liver sections from the mice carrying targeted gene KO and fed with 3 months of WD. A. H&E staining of liver sections from control KO mice. The same 15x images for *GFP* and *LacZ* are also shown in Figure 5J; they are included here for the purposes of comparison. B. H&E staining of liver sections from NASH gene KO mice. The same 15x images for *Irs1*, *Srebf1* and *Dgat2* are also shown in Figure 5J. C. H&E staining of liver sections from transcription factor KO mice. The same 15x images for *Tbx3* and *Bcl6* are also shown in Figure 5J. D. H&E staining of liver sections from epigenetic factor KO mice. The same 15x images for *Smyd2* are also shown in Figure 5J.

## Notes

### Competing Interest Statement

Y.H. consults for Helio Genomics, Espervita Therapeutics, and Roche Diagnostics. Y.H. is a shareholder of Alentis Therapeutics and Espervita Therapeutics. M.H. consults for Spliceor, is on the speakers panel for Boston Scientific, and has research support from Pfizer and AstraZeneca. H.Z. consults for Alnylam Pharmaceuticals, Jumble Therapeutics, and Chroma Medicines, and serves on the SAB of Ubiquitix. H.Z. has research support from Chroma Medicines. H.Z. owns stock in Ionis and Madrigal Pharmaceuticals. M.H., H.Z., and P.C. consult for FL86 and Flagship Pioneering. M.H. and P.C. are co-inventors on patents on somatic mutants in liver disease, including ACVR2A and GPAM. Z.W., H.Z., and L.L. are co-inventors on patents on GPAM, TBX3, and SMYD2 siRNAs.

